# A lateral linker histone binding mode scaffolds dinucleosome stacking in chromatin fibers

**DOI:** 10.64898/2026.03.29.715057

**Authors:** Youchao Wang, Shuoming Yang, Jiachen Wang, Xiaolong Wu, Dejian Zhou, Peng Liu, Ran Zha, Junzhe Sun, Jinru Zhang, Jinzhong Lin, Huabin Zhou, Jianhua Gan, Aiwu Dong

**Author notes:** Corresponding author. (H.Z.); (J.G.); (A.D.).

## Abstract

Linker histones are essential for chromatin compaction, yet how they contribute to higher-order fiber assembly remains poorly understood. Here, we determined cryo-electron microscopy structures of Arabidopsis dodeca-nucleosome fibers containing distinct H2A/H3 variants and linker histone H1.3, revealing a noncanonical binding mode that a laterally positioned H1.3 connects the acidic patch of one nucleosome and the DNA of the neighboring nucleosome, thereby scaffolding dinucleosomes into two-start chromatin fibers. This noncanonical binding mode is structurally conserved when H1.3 is replaced by *Gallus gallus* H5. Furthermore, incorporation of H2A.W and H3.3 further induces back-to-back fiber dimerization. Cryo-electron tomography and *in vivo* cross-linking mass spectrometry analyses support the physiological relevance of H1 lateral engagement. Our findings establish that linker histones act as active architectural scaffolds in higher-order chromatin organization.

## Main Text

By organizing DNA into nucleosomes, 30-nm fibers and higher-order assemblies, chromatin achieves a dynamic balance between compaction and accessibility, thereby regulating DNA replication, transcription, and repair (*1, 2*). The incorporation of histone variants influences nucleosome stability, inter-nucleosomal interactions, recruitment of chromatin-associated factors, and chromatin structures (*3, 4*). High-resolution structures of mononucleosomes containing distinct histone variants from different species have been determined (*5–11*). Extensive structural studies of reconstituted chromatin fibers have revealed how nucleosomes assemble into higher-order 30-nm fibers and suggested diverse interaction modes that mediate nucleosome stacking, involving histone tails, linker histones, and linker DNA geometry (*12–18*). However, whether there are new configurations for 30-nm fibers, or whether alternative assembly modes exist, remains elusive.

Central to this question is the linker histone. In all structurally characterized chromatin fibers to date, linker histones bind at or near the nucleosomal dyad, where they engage linker DNA to constrain its trajectory and promote fiber compaction (*12–14, 16*). Yet biochemical and computational studies have suggested that linker histones can adopt noncanonical binding configurations beyond the nucleosomal dyad, including positions that may approach the H2A–H2B acidic patch, a nucleosomal surface widely exploited by diverse chromatin regulators (*19, 20*). Whether such alternative binding modes are realized within chromatin fibers, and what architectural roles they might play, are still unknown.

Plants provide a compelling system to address the above issues. As sessile organisms, plants rely on rapid and reversible chromatin-based regulation to adapt to fluctuating environmental conditions. Arabidopsis encodes functionally divergent linker histone variants, including H1.3, a stress-inducible isoform implicated in environmental stress responses, chromatin remodeling, DNA methylation, and transcriptional regulation (*21–24*). Plants possess lineage-specific histone variants, such as heterochromatin-enriched H2A.W (*25–28*), whose contribution to higher-order chromatin structure has not been explored. Here, we reconstituted plant chromatin fibers *in vitro* using Arabidopsis histones and determined their structures by cryo-electron microscopy (cryo-EM) single-particle analysis. These structures reveal previously unrecognized features of chromatin fiber organization and provide new structural insights into how linker histones contribute to higher-order chromatin architecture.

### Arabidopsis chromatin assembles into a dinucleosome-based chromatin fiber structure

Do plant core histones and histone variants assemble distinct higher-order chromatin structures? To address this question, we reconstituted chromatin fibers *in vitro* using Arabidopsis histones (H2A.Z, H2B, H3.1, H4 and linker histone H1.3) and 12 tandem repeats of 177 bp (12 × 177 bp) DNA. Structures of the mononucleosomes used in fiber reconstitution, including those containing H2A/H2A.Z/H2A.W and H3.1/H3.3, are overall similar (*11*) (fig. S1-S4 and table S1). The structure of the dodeca-nucleosomal Arabidopsis chromatin fiber was resolved by cryo-EM single-particle analysis at ∼10 Å resolution, whereas focused refinement of a single nucleosome yielded a local reconstruction at 2.87 Å resolution (fig. S5 and table S2).

Although the Arabidopsis chromatin fiber followed the classic two-start zig-zag mode, it displayed a different structure from the previously reported H1- or H5-bound chromatin fibers (Fig. 1, fig. S6, and tables S3 and S4) (*12–14, 16*). First, instead of tetranucleosomal units within the H5-bound chromatin (H5-chromatin) fiber, the left-handed double-helical structure of Arabidopsis chromatin fiber was formed by zig-zag dinucleosomal units (Fig. 1A-C and fig. S6A-C). Second, compared with the H5-chromatin fiber structure, the dinucleosome distance (between nucleosomes N and N+1) was reduced, whereas the opening angles (between nucleosomes N and N+1) along the nucleosome array were increased in the Arabidopsis chromatin fiber (Fig. 1D and 1E, fig. S6D–H, and table S4). Third, the stacking distance between the adjacent nucleosomes (N and N+2) of the Arabidopsis chromatin fiber was significantly longer than that of H5-chromatin fiber (Fig. 1F, 1G and 1H, and table S3). Fourth, the disc planes of nucleosomes N and N+2 in the Arabidopsis chromatin fiber were more parallel than those in the H5-chromatin fiber (Fig. 1E and 1F, and fig. S6D-F, and table S3). Taken together, the above features of Arabidopsis chromatin fiber produce a more open and ladder-like stacking geometry, which is distinct from the known tetranucleosomal fiber architectures.

**Fig. 1.**
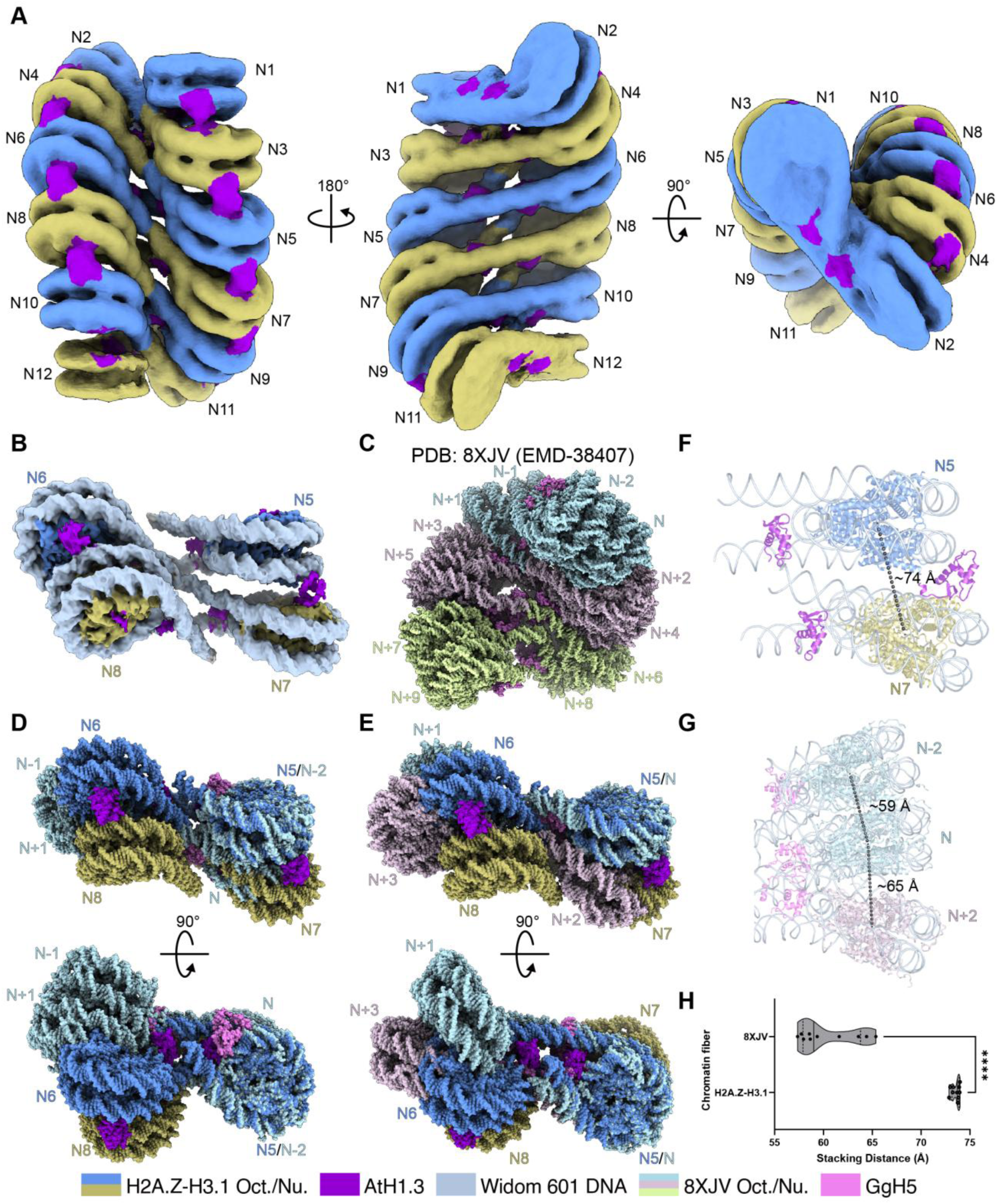
Structural basis of chromatin fibers reconstituted with Arabidopsis histones H1.3, H2A.Z, H2B, H3.1, and H4. (**A**) Cryo-EM density map of a reconstituted chromatin fiber containing twelve nucleosomes (N1–N12) assembled with Arabidopsis H1.3 (AtH1.3), H2A.Z, H2B, H3.1, and H4. Adjacent dinucleosome units are colored differently, with linker histone H1.3 highlighted in dark violet. The map has been deposited in the Electron Microscopy Data Bank (EMDB) under accession number EMD-67700. Oct./Nu. indicates octamer/nucleosome. (**B**) Focused cryo-EM map of four nucleosomes within the H2A.Z-containing chromatin fiber (A). DNA, linker histone H1.3, and histone octamers are colored distinctly. The map has been deposited in the EMDB under accession number EMD-67701. (**C**) The atomic model of the published H5-chromatin fiber (PDB: 8XJV; EMD-38407) containing twelve nucleosomes (N-2–N+9), in which each tetranucleosomal unit is shown in a distinct color. Linker histone H5 is highlighted in violet. (**D**) Structural alignment of the Arabidopsis H2A.Z-containing tetranucleosome (cornflower blue, dark khaki and dark violet) with the tetranucleosomal unit of PDB 8XJV (light blue and violet). (**E**) Structural alignment of the Arabidopsis H2A.Z-containing tetranucleosome (cornflower blue, dark khaki and dark violet) with a segment spanning two adjacent tetranucleosomal units of PDB 8XJV, containing the last two nucleosomes of the first unit and the first two nucleosomes of the second unit (light blue, thistle and violet). (**F**) Distances between adjacent nucleosomes measured from an atomic model built into a reconstructed dinucleosome composite cryo-EM map of the Arabidopsis H2A.Z-containing chromatin fiber. The composite map has been deposited in the EMDB under accession number EMD-67706, and the two focused maps used for composite map generation have been deposited under accession numbers EMD-67702 and EMD-67703. The atomic coordinates have been deposited in the Protein Data Bank (PDB) under accession number 21IV. (**G**) Distances between adjacent nucleosomes within a single tetranucleosomal unit of PDB 8XJV, and between neighboring nucleosomes across adjacent tetranucleosomal units. (**H**) Violin plot comparing nucleosome stacking distances measured for the twelve nucleosomes in the Arabidopsis H2A.Z-containing chromatin fiber shown in panel (A) and those in the PDB 8XJV chromatin fiber. Note that the distances measured here are derived from the chromatin fiber map and may differ from those obtained from the atomic model built into the dinucleosome composite cryo-EM map, as shown in panel (F). Statistical significance was assessed using the Mann-Whitney test (P < 0.0001).

### Histone H1.3 bridges adjacent nucleosomes through a noncanonical lateral binding mode

In the Arabidopsis fiber structure, each nucleosome interacted with two H1.3 molecules, here referred to as H1.3_A and H1.3_B (Fig. 2A and 2B). H1.3_A occupied the canonical on-dyad site, where its globular domain (GD) formed asymmetric interactions with the nucleosomal DNA, consistent with the previously reported H1 and H5 structures in the animal chromatin fibers (Fig. 2B, fig. S6I and S6J) (*29, 30*). By contrast, H1.3_B adopted a lateral position on the surface of nucleosomes and mediated a previously unobserved nucleosome-bridging interaction. This bridging was achieved through the two domains of H1.3_B: its C-terminal domain (CTD) anchored to nucleosome N, while its GD spanned and contacted the DNA of the neighboring nucleosome N+2 (Fig. 2B).

**Fig. 2.**
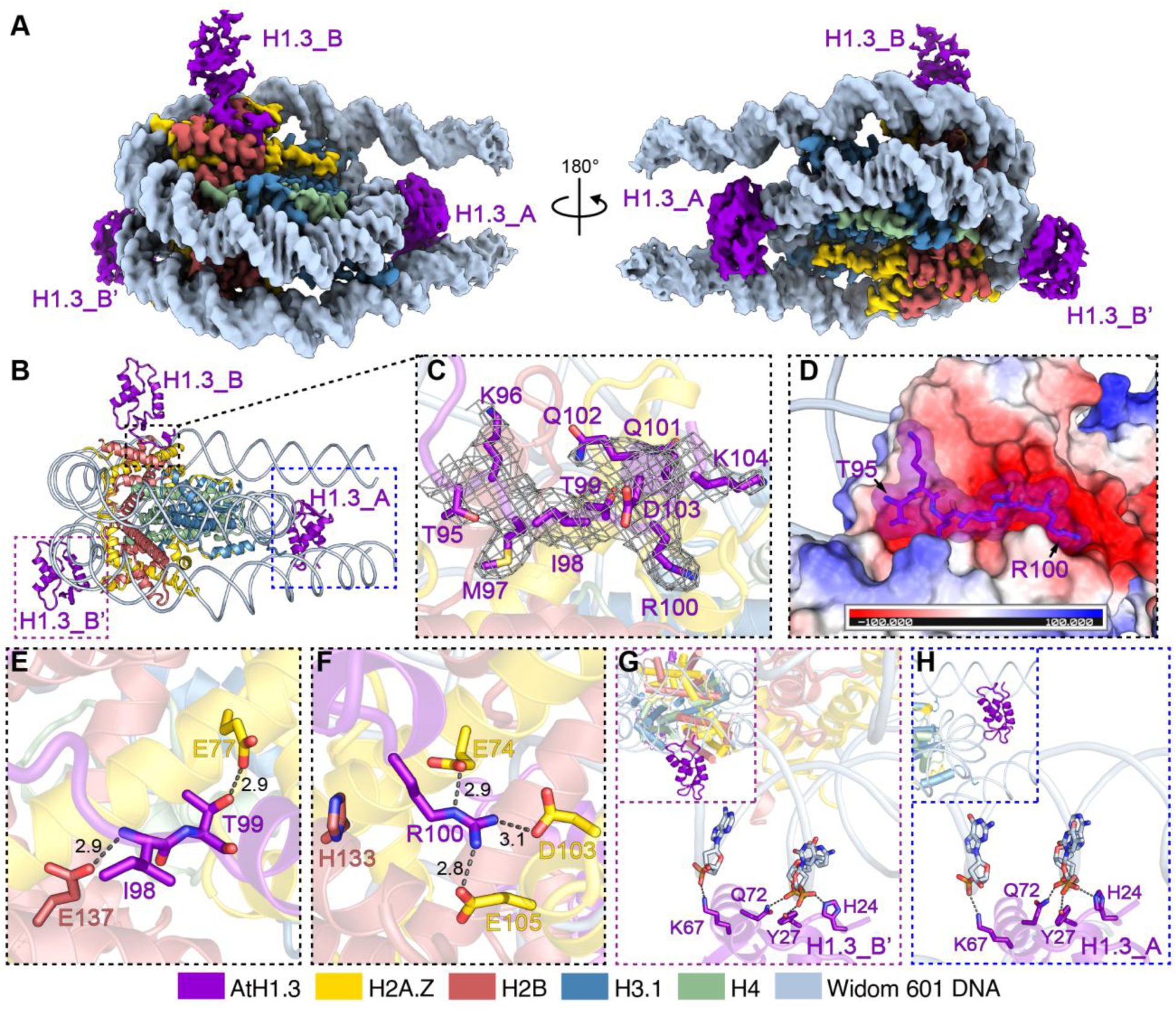
Molecular basis of linker histone H1.3-mediated nucleosome stacking. (**A**) Focused cryo-EM map showing a single nucleosome within the Arabidopsis H2A.Z-containing chromatin fiber associated with two H1.3 molecules (H1.3_A and H1.3_B). Density corresponding to H1.3_B in the neighboring nucleosome is labeled as H1.3_B’. Each chain is colored distinctly. The map has been deposited in the EMDB under accession number EMD-67705. (**B**) Atomic model corresponding to (A), illustrating dual H1.3 engagement in a single nucleosome. The atomic coordinates have been deposited in the PDB under accession number 21IU. (**C**) Cryo-EM density corresponding to the residues T95-K104 within the C-terminal of H1.3. (**D**) Position of the C-terminal basic cluster of H1.3 (residues T95-R100) in the acidic patch of a nucleosome. (**E**) Hydrogen-bond interactions between H1.3 Ile98 and H2B Glu137, and between H1.3 Thr99 and H2A.Z Glu77. (**F**) Hydrogen-bond interactions between H1.3 Arg100 and H2A.Z residues Glu74, Asp103, and Glu105, as well as a cation-π interaction with H2B His133. (**G**, **H**) The laterally bound H1.3 (G) and the H1.3 at the on-dyad site (H) engage nucleosomal DNA through an identical binding interface. Selected hydrogen bonds between H1.3 side chains and the backbone phosphate groups of the nucleosomal DNA are shown. The inset (upper left) indicates the position of the interaction within the overall structure.

The N-terminal domain (NTD) of H1.3_B was disordered, whereas a short segment of its CTD (residues 95–104) was well resolved, supported by clear electron density (Fig. 2C). Atomic modeling revealed that this CTD segment inserted directly into the nucleosomal acidic patch (Fig. 2D), a negatively charged surface formed primarily by residues from H2A.Z and H2B. Residues from both histones contributed to stabilizing the H1.3_B CTD through electrostatic and hydrogen-bonding interactions. As depicted in Fig. 2E, the backbone amide nitrogen of H1.3_B Ile98 formed hydrogen bonds with the side chain of H2B Glu137, while H1.3_B Thr99 engaged H2A.Z Glu77 through hydrogen bonding. Moreover, H1.3_B Arg100 established additional contacts by forming hydrogen bonds with H2A.Z Glu74, Asp103, and Glu105, as well as a cation-π interaction with H2B His133 (Fig. 2F). Notably, H1.3_B Arg100 is positively charged, whereas H2A.Z Glu74, Asp103, and Glu105 are all negatively charged. Consistent with their high sequence conservation among Arabidopsis H2A variants (fig. S7A), these acidic residues may also contribute to H1.3_B interactions in chromatin fibers containing H2A or H2A.W.

No stable interaction was observed between the GD and CTD of H1.3_B. Consequently, the GD adopted a flexible conformation and variable orientation, as indicated by its weaker density maps (Fig. 2A). In contrast to the CTD, which anchored H1.3_B to the acidic patch of nucleosome N, the GD extended laterally to engage the DNA of the adjacent nucleosome N+2. As depicted in Fig. 2G, His24 and Tyr27, located at the N-terminal α1 helix of the GD, formed hydrogen bonds with the backbone phosphate groups of the nucleosomal DNA. The α3 helix of GD inserted into the minor groove of the N+2 nucleosomal DNA, and the side chains of Lys67 and Gln72 extended toward the phosphate backbone to form additional hydrogen bonds (Fig. 2G). The DNA-interacting interface within H1.3_B was the same as that within H1.3_A (Fig. 2H). The four residues of H1.3 responsive for interacting with DNA are also conserved in Arabidopsis H1.1 and H1.2 (fig. S7B), suggesting that these H1 variants may likewise assembly similar chromatin fibers as H1.3.

To examine whether the observed Arabidopsis chromatin fiber architecture is specific to H2A.Z, we reconstituted chromatin fibers containing canonical H2A or the plant-specific variant H2A.W and determined their structures (figs. S8 and S9, tables S5 and S6). Although fibers assembled with H2A or H2A.W exhibited increased conformational heterogeneity and greater flexibility, they both adopted a nucleosome organization mode similar as that of the H2A.Z-containing chromatin (H2A.Z-chromatin) fiber (fig. S10). H2A.Z and H2A.W share high sequence similarity with H2A within their core regions, whereas their N- and C-terminal tails differ substantially (fig. S7A). Consistent with these observations, our cryo-EM structures showed that the linker histone H1.3 mediated nucleosome stacking primarily through interactions with the conserved histone core region rather than the disordered histone tails.

Taken together, these findings indicate that H1.3 mediates nucleosome stacking through a division of labor between its CTD, which engages the acidic patch, and its GD, which stabilizes interactions with the neighboring nucleosomal DNA. Furthermore, this noncanonical lateral binding mode mediated by Arabidopsis H1.3 is independent of sequence variation within the N-and C-terminal regions of H2A variants.

### The lateral bridging mode is conserved when replacing Arabidopsis H1.3 by *Gallus gallus* H5

To test whether the H1.3-mediated stacking mode represents a more general feature of linker histones, we reconstituted chromatin fibers containing *Gallus gallus* linker histone H5 and Arabidopsis H2A.Z, H2B, H3.1, H4 and 12 × 177 bp DNA. The structure of H5-H2A.Z-chromatin fiber was determined by cryo-EM at 3.92 Å resolution (fig. S11 and table S7). Like the H1.3-H2A.Z-chromatin, the H5-H2A.Z-chromatin assembled into a two-start double-helical chromatin fiber (Fig. 3A and 3B, fig. S11C). Notably, the distance between nucleosomes N and N+2 in the H5-H2A.Z-chromatin was comparable to that observed in the H1.3-H2A.Z-chromatin fiber (Fig. 1F and 3C), indicating that the lateral spacing between stacked nucleosomes is largely preserved.

**Fig. 3.**
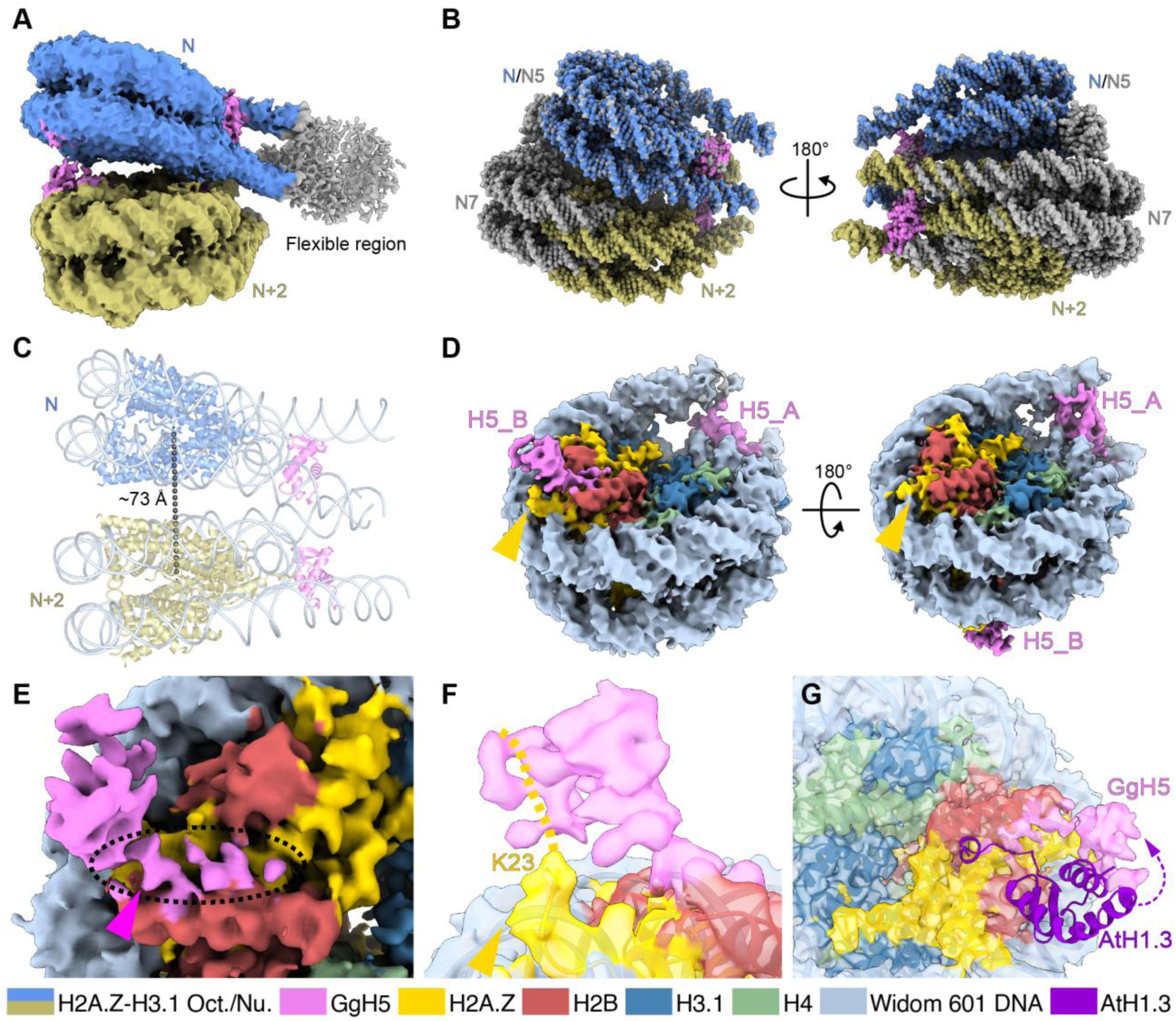
Structural organization of chromatin fibers assembled with *Gallus gallus* H5. (**A**) Overall cryo-EM density map of the chromatin fiber assembled with Arabidopsis H2A.Z, H2B, H3.1, H4 and *Gallus gallus* H5 (GgH5). The nucleosomes are colored distinctly. The map has been deposited in the EMDB under accession number EMD-67719. (**B**) Structural alignment of the H5-containing dinucleosome (PDB: 21IY, cornflower blue, dark khaki and violet) with the H1.3-containing dinucleosome (PDB: 21IV, gray). (**C**) Distances between adjacent nucleosomes measured from an atomic model built into three cryo-EM maps of the H5-H2A.Z-containing chromatin fiber. The overall chromatin fiber map shown in (A) (EMD-67719), and the two focused maps used for model refinement have been deposited under accession numbers EMD-67717 and EMD-67718. The atomic coordinates have been deposited in the PDB under accession number 21IY. (**D**) Focused cryo-EM map showing a single nucleosome associated with an upper-positioned H5 molecule. Each chain is colored distinctly. An orange arrow indicates additional density adjacent to the N terminus of H2A.Z. The map has been deposited in the EMDB under accession number EMD-67720. (**E**) Cryo-EM density corresponding to H5 bound to the nucleosome, showing discontinuous density for the C-terminal region of H5. (**F**) Atomic model of the H2A.Z-containing nucleosome fitted into the density shown in (D). A dashed line indicates the potential extension direction of the N-terminal tail of H2A.Z. The atomic coordinates have been deposited in the PDB under accession number 21IZ. (**G**) Atomic model of the H2A.Z-containing nucleosome bound by an upper-positioned Arabidopsis H1.3 (PDB: 21IU) fitted into the density shown in (D), illustrating a substantial positional offset between H5 and H1.3.

In the H5-H2A.Z-chromatin fiber, each nucleosome interacted with two H5 molecules, referred to as H5_A and H5_B (Fig. 3D). H5_A bound at the on-dyad site, adopting an orientation and interaction mode similar as H1.3_A in the H1.3-H2A.Z-chromatin structure. Like H1.3_B, H5_B occupied a lateral position on one nucleosome and extended to the adjacent nucleosome, consistent with a bridging role (Fig. 3A and 3D). In this structure, extended cryo-EM densities were observed adjacent to the nucleosomal acidic patch, likely corresponding to a segment of the H5 CTD (Fig. 3E). Although the density was insufficiently resolved to assign individual residues, likely reflecting conformational heterogeneity at this interface, it supported that the H5 CTD engaged in the acidic patch, and the density corresponding to the N-terminal region of H2A.Z was positioned in proximity to H5 at this interface (Fig. 3D and 3F), suggesting that the N-terminal tail of H2A.Z may contribute to stabilizing H5 binding.

Despite this shared structural framework, the two types of fibers differed in detail. H5-H2A.Z-chromatin fibers adopted a more twisted overall conformation, and the relative orientation of H5_B differed from that of H1.3_B (Fig. 3A and 3G, and fig. S11C). These differences are likely due to sequence variations in the CTD: although the CTDs of H1.3 and H5 are both enriched in positively charged residues that can engage in electrostatic interactions with acidic patches, their amino acid sequences are not conserved (fig. S7C). The close proximity between the H5 GD and the neighboring nucleosomes further supported the possibility of direct nucleosome-to-nucleosome stabilization. This suggested a model in which GD-DNA interactions are conserved, mediated by common structural features of the GD, whereas CTD-acidic patch interactions are adaptable, capable of tolerating sequence differences while maintaining overall bridging function.

Taken together, these observations indicate that linker histone–mediated nucleosome stacking represents a conserved architectural principle rather than a plant-specific specialization. Although different linker histones may vary in sequence and conformational details, the ability to link nucleosomes by coordinating the binding of DNA and acidic patches appears to be evolutionarily conserved.

### The incorporation of H3.3 and H2A.W variants causes back-to-back dimerization of chromatin fibers

Besides H2A variants, H3.3 is another histone variant that is very important in eukaryotes, distributed across both euchromatin and heterochromatin. H3.3 is traditionally associated with active transcription, however, it also functions in maintaining heterochromatin integrity and telomere stability (*31*). Compared to the canonical histone H3.1, Arabidopsis H3.3 contains four substitutions: A32T at the N-terminus and F42Y, S88H, and A91L within the core region (fig. S7D). To investigate whether variation of H3 impacts chromatin fiber architecture, we reconstituted chromatin fibers by replacing H3.1 with H3.3. Due to pronounced conformational heterogeneity, we were unable to obtain stable and well-resolved chromatin fiber structures containing H3.3 together with H2A.Z or canonical H2A. In contrast, chromatin fibers containing H2A.W, H3.3 and H1.3 (H3.3-H2A.W-chromatin) yielded structurally interpretable reconstructions (fig. S12), which is consistent with our observation that nucleosomes containing H3.3 and H2A.W exhibited markedly higher structural stability and homogeneity in both representative micrographs and 3D classifications compared with H2A- or H2A.Z-containing nucleosomes (fig. S1, S2 and S3). H3.3-H2A.W-chromatin fibers were therefore selected for further analysis. Interestingly, two different types of structures were observed, with ∼93% of particles belonging to monomeric fibers and ∼7% to dimeric fibers (fig. S12, tables S3, S4 and S8). The assembly and overall conformation of the monomeric fibers were similar as those of H3.1-containing chromatin fibers (Fig. 4A and 4B), whereas the structure of the H3.3-H2A.W dimeric fibers was markedly different from all known chromatin fibers, with two chromatin fibers arranged in a back-to-back configuration (Fig. 4C and 4D). In both types of H3.3-H2A.W-chromatin structures, the linker DNAs of the nucleosomes bent on one side while remaining relatively straight on the opposite side, and chromatin dimerization occurred on the straight linker DNA side. Structural analysis revealed that the stacking distances between nucleosomes N and N+2 were similar in the two types of H3.3-H2A.W-chromatin fibers but significantly increased compared with that in H2A.Z-H3.1-chromatin fibers (Fig. 4E and 4F, and table S3). Moreover, the H3.3-H2A.W dimeric fibers exhibited significantly different distances and angles compared with H2A.Z-H3.1-chromatin fibers, while showing no significant differences relative to H3.3-H2A.W monomeric fibers. (Fig. 4E and 4F, fig. S13 and tables S3 and S4). The stacking angles between nucleosomes N and N+2 and rotation angles between neighboring dinucleosomes (N-1–N and N+1–N+2) were relatively smaller in the H3.3-H2A.W dimeric fibers, which created a flatter surface and possibly facilitated dimerization between the monomeric fibers (Fig. 4F, fig. S13E, and table S4). Although the overall length of the two types of H3.3-H2A.W-chromatin fibers was comparable, the width of the dimeric fiber (∼300 Å) was greater than that of the monomeric fiber (∼230 Å) (fig. S13A and S13B). At the current resolution (∼15 Å), it is difficult to identify the specific residues mediating back-to-back dimerization in the dimeric H3.3-H2A.W-chromatin fibers. Nevertheless, structural analysis indicated that the N-terminus of H3.3 extended toward the dimerization interface. Notably, the N-terminal region contains the A32T substitution, which may enhance interactions either between the N-termini of H3.3 molecules from opposing fibers or between the H3.3 N-terminus and other histone regions, such as the C-terminus of H2A.W (Fig. 4G-I). Such interactions could contribute to stabilization of this higher-order chromatin assembly.

**Fig. 4.**
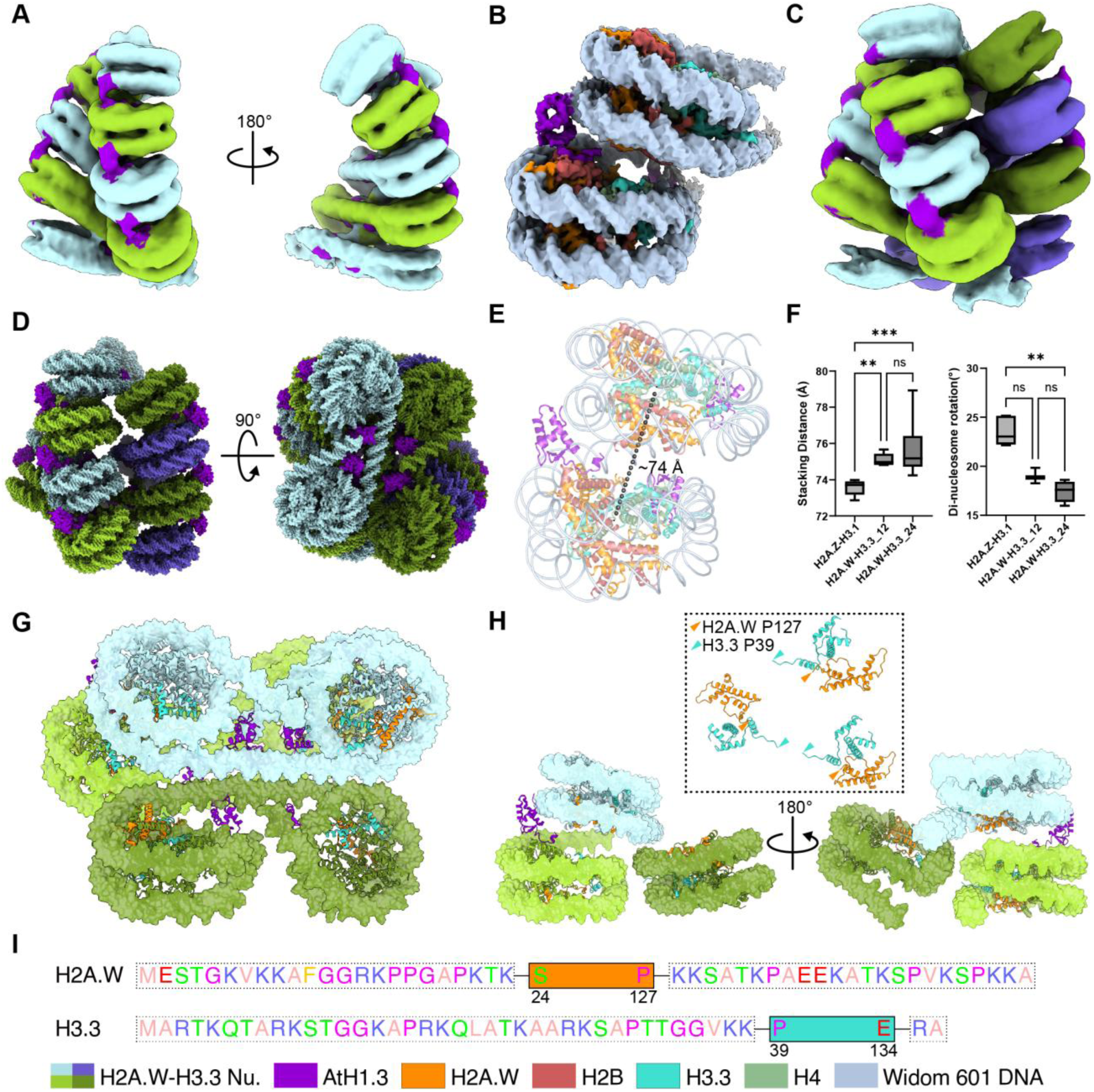
Structure of Arabidopsis chromatin fibers containing histones variants H2A.W and H3.3. (**A**) Cryo-EM density map of a reconstituted chromatin fiber assembled with Arabidopsis H1.3, H2A.W, H2B, H3.3, and H4. Adjacent dinucleosome units are colored differently, with linker histone H1.3 highlighted in dark violet. The map has been deposited in the EMDB under accession number EMD-68924. Nu. indicates nucleosome. (**B**) Focused cryo-EM map of stacking dinucleosomes within the H2A.W-H3.3-containing chromatin fiber. Each chain is colored distinctly. The map has been deposited in the EMDB under accession number EMD-67723. (**C**) A specific classification revealed a higher-order chromatin organization formed by back-to-back fiber dimerization, with eight nucleosomes clearly visible in each fiber. The map has been deposited in the EMDB under accession number EMD-67724. (**D**) Overall interaction model of the two chromatin fibers shown in panel (C), illustrated by atomic models. (**E**) Distances between adjacent nucleosomes measured from an atomic model built into three cryo-EM maps of the Arabidopsis H2A.W-H3.3-containing chromatin fiber. The overall chromatin fiber map shown in (B) (EMD-67723), and the two focused maps used for model refinement have been deposited under accession numbers EMD-67721 and EMD-67722. The atomic coordinates have been deposited in the PDB under accession number 21JA. (**F**) Box plots comparing nucleosome stacking distances (left) and rotation angles between adjacent dinucleosome units (right) measured for the Arabidopsis H2A.Z-H3.1-containing chromatin fiber (H2A.Z-H3.1), the H2A.W-H3.3-containing chromatin fiber (H2A.W-H3.3_12), and the H2A.W-H3.3-containing fiber dimerization (H2A.W-H3.3_24). Distances were measured directly from chromatin fiber maps and may therefore differ from values obtained from atomic models built into focused dinucleosome cryo-EM maps, as shown in panel (E). Statistical significance was assessed using the Kruskal-Wallis test followed by Dunn’s multiple comparisons test. Asterisks indicate levels of significance (*P < 0.05; **P < 0.01; ***P < 0.001); ns, not significant. (**G**) Close-up view of the interaction interface between two chromatin fibers, with four nucleosomes shown on one side and two nucleosomes on the other. DNA is displayed as a surface representation, whereas histones are shown in cartoon representation. Histones H1.3, H2A.W, and H3.3 are colored differently for distinction. (**H**) A more detailed view of the interactions shown in panel (G), with two stacked nucleosomes on one side and a single nucleosome on the other. The dashed box highlights the relative positions of H2A.W and H3.3 histones from three nucleosomes that may participate in the interaction. Dark orange arrows indicate residues P127 (residue 128 and downstream residues of H2A.W that were not resolved), while cyan arrows indicate residues P39 (residue 38 and upstream residues of H3.3 that were not resolved). (**I**) Amino acid sequences of the unresolved intrinsically disordered regions of H2A.W (top) and H3.3 (bottom) highlighted in the dashed box. Amino acids are shown in single-letter code and colored according to their physicochemical properties.

### *In vivo* cross-linking and tomographic analyses support lateral linker histone engagement

Since both Arabidopsis H1.3 and *Gallus gallus* H5 achieved the formation of noncanonical 30-nm chromatin fibers, we propose that linker histone–mediated nucleosome stacking represents a general assembly principle *in vitro*. To verify whether linker histones also mediate such nucleosome stacking *in vivo*, we first analyzed the reported cross-linking mass spectrometry (XL-MS) data identified from soybean and human nuclei (*32, 33*), focusing on interactions between linker histones and core histones. Several cross-links support the H1.3-mediated nucleosome stacking model, in which H1 anchors to the nucleosomal DNA via its GD while simultaneously capturing the acidic patch of the adjacent nucleosome through its flexible C-terminal tail (Fig. 5A-E, fig. S14 and table S9).

**Fig. 5.**
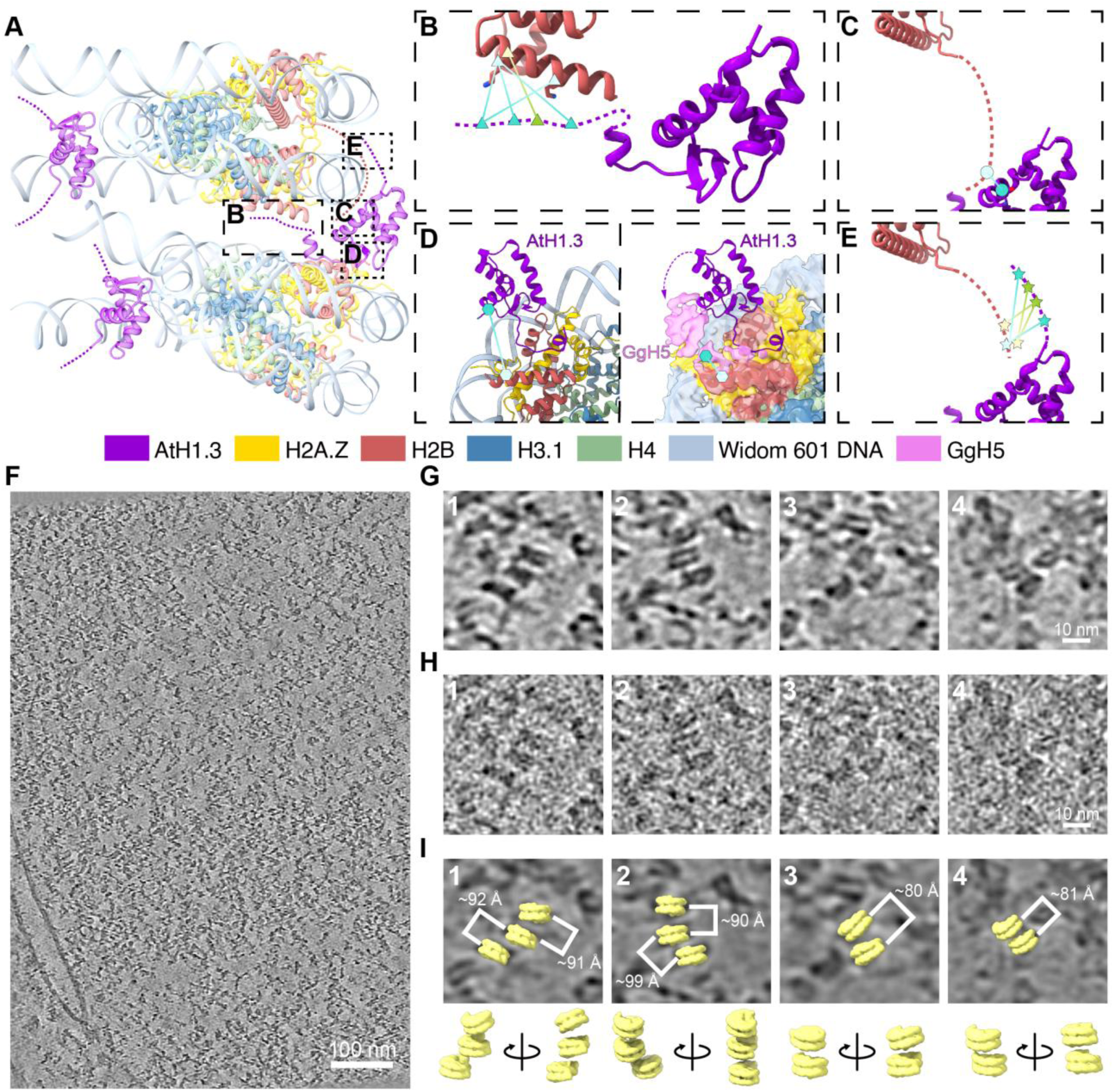
*In vivo* evidence supporting linker histone-mediated nucleosome stacking. **(A)** Atomic model of a stacked dinucleosome configuration (PDB: 21IV). Each chain is colored distinctly. Dashed lines indicate the possible extension of the N- and C-terminal regions of histone H1.3 (dark violet) and H2B (Indian red). The boxed regions are enlarged into (B–E). **(B–E)** Cross-linking mass spectrometry (XL-MS) data of soybean and human nuclei mapped back to (A). Cross-links identified in soybean and human nuclei are shown in different colors, and distinct symbols denote different cross-linking regions. Panel (D) additionally includes the cryo-EM density of the H5-H2A.Z-containing chromatin fiber, illustrating that H5 is positioned closer to H2B in this region compared with Arabidopsis H1.3. (**F**) Denoised cryo-electron tomographic slice of a nuclear region in a rice protoplast (slice 126). The nuclear envelope, nuclear pores, and densely packed chromatin within the nucleus are visible. Both the original tomogram reconstructed using Warp and the corresponding denoised tomogram have been deposited in the Electron Microscopy Data Bank (EMDB) under accession number EMD-67852. (**G**) Enlarged views of four representative nucleosome-stacking regions from the denoised tomogram shown in panel (F). The four regions are numerically labeled, and their corresponding IMOD coordinates are: 1 (567, 205, 117), 2 (532, 99, 126), 3 (150, 747, 135), 4 (276, 869, 155). (**H**) Corresponding views of the same four regions shown in panel (G), extracted from the original, non-denoised tomogram reconstructed using Warp. The numbering matches that in panel (G). (**I**) Three-dimensional visualization of the nucleosome-stacking regions shown in panel (G) by fitting a low-resolution mononucleosome density map into the tomogram at the corresponding coordinates. The fitted nucleosome densities (top) and their relative spatial arrangements displayed in 3D (bottom) are shown for each region, with numbering corresponding to panel (G). Euclidean distances between the centers of neighboring nucleosome particles are indicated. Because nucleosomes are disk-shaped and may adopt tilted orientations, the minimal distances between adjacent DNA surfaces can be smaller than the center-to-center distances shown. This analysis illustrates the relative positioning and stacking geometry of nucleosomes within the native chromatin context.

The identified cross-links can be grouped into four categories. (1) Multiple cross-links connected the C-terminal region of H1 to two fixed positions within the C-terminal α4 helix of H2B (Fig. 5A and 5B; fig. S14; table S9). In our atomic model, the C-terminal region of H1 could easily extend to these positions, and the observed cross-links are consistent with a highly dynamic H1 C-terminus. (2) Cross-links were observed between the intrinsically disordered N-terminal region of H2B and helix α3 of H1 (Fig. 5A and 5C; fig. S14; table S9). The N-terminus of H2B was sufficiently flexible to reach this region, consistent with lateral binding of H1 on the nucleosome surface. (3) In human nuclei, cross-links were detected between the regions spanning helices α1 and α2 of linker histone H1.5 and the C-terminal helix α4 of histone H2B (Fig. 5A and 5D; fig. S14; table S9). In the H1.3-H2A.Z-chromatin fiber structure, the corresponding positions were spatially separated, whereas in the H5-H2A.Z-chromatin fiber structure, these sites were close to each other (Fig. 3G and Fig. 5D). This observation suggests that human H1.5 may adopt a nucleosome-stacking configuration similar to that of *Gallus gallus* H5 in the H5-H2A.Z-chromatin fiber, which is also consistent with the recent study reporting the interaction between human H1.5 and centromeric protein A (CENP-A) mononucleosomes (*20*). (4) Cross-links were detected between the N-terminal regions of H1 and H2B (Fig. 5A and 5E; fig. S14; table S9). In the lateral H1-binding configuration observed in our structures, the N-terminus of H1 is located next to the N-terminus of H2B of the adjacent upper nucleosome, providing a structural basis for these interactions.

To examine whether linker histones mediate the stacking of neighboring nucleosomes *in vivo*, we performed cryo-electron tomography (cryo-ET) of lamellae containing nuclear regions from rice protoplasts. Rice histones and histone variants are highly conserved with those of Arabidopsis (fig. S15) (*34*). In addition, the rice genome is substantially larger and contains a higher proportion of repetitive and heterochromatic sequences than the Arabidopsis genome, which may help in the *in situ* visualization of higher-level chromatin organization. Tomographic reconstruction revealed regions of highly condensed chromatin, where nucleosomes were stacked in a stepwise manner (Fig. 5F-I).

The spacing and geometry of nucleosomes (∼80–92 Å) in rice protoplasts were close to the dinucleosome stacking distance (∼74–76 Å) observed in our Arabidopsis chromatin fibers, while notably deviating from the more compact stacking (∼57–64 Å) reported for animal H1/H5-containing 30-nm chromatin fibers (EMDB: EMD-2600; PDB: 8XJV) (Fig. 5I and table S3). It should be noted that because nucleosomes are disk-shaped and often adopt tilted orientations, the minimal distances between adjacent DNA surfaces can be smaller than the center-to-center distances shown. Furthermore, variations in the C-terminal tail length among H1 isoforms may contribute to the increased stacking distances mediated by this lateral binding mode (fig. S7B). Although the resolution of cryo-ET (∼30–40 Å) is insufficient to identify linker histones at the stacking interface, the observed geometry was consistent with the role of linker histone H1 in stabilizing such stacked nucleosome configuration. This is further supported by recent *in situ* structural studies of intact human nuclei (*35, 36*), which consistently show that the majority of dinucleosome stacking distances in interphase chromatin exceed 70 Å. Collectively, our cryo-ET observations, together with previously reported cryo-ET studies (*18, 35–40*) and XL-MS constraints from different species, support the existence of nucleosome stacking configurations mediated by linker histones *in vivo* and are consistent with a model in which linker histones connect adjacent nucleosomes through a noncanonical lateral binding mode.

## Discussion

The classical model of linker histone function portrays H1 as a monovalent organizer of linker DNA, binding the nucleosome dyad, constraining linker DNA exit angles, and thereby promoting fiber compaction indirectly (*12–14, 16, 41*). Our structures demand a fundamental revision of this view. We show that H1.3 simultaneously occupies two functionally distinct positions on each nucleosome: a canonical dyad site where it constrains linker DNA geometry, and a previously undescribed lateral position from which it directly bridges the adjacent nucleosomes by engaging its acidic patch and DNA in trans. This division of labor transforms H1.3 from a passive architectural element into an active inter-nucleosomal scaffold, with broad implications for how linker histone sequence diversity is mechanistically coupled to chromatin structural diversity.

To determine whether the lateral bridging mechanism is a plant-specific specialization or a general feature of linker histone function, we replaced Arabidopsis H1.3 with *Gallus gallus* H5. Despite substantial CTD sequence divergence between the two proteins, both linker histones support a structurally similar lateral bridging configuration, with the CTD engaging the nucleosomal acidic patch and the GD contacting neighboring nucleosomal DNA. This convergence indicates that the critical determinant of lateral bridging capacity is electrostatic rather than sequence-specific: a sufficiently basic C-terminal patch with appropriate conformational flexibility may be sufficient for acidic patch engagement, irrespective of primary sequence. Unlike core histones, which are highly conserved, linker histones exhibit substantial sequence divergence and functional diversification across eukaryotes (*42, 43*). The independent acquisition of lateral bridging capacity in phylogenetically distant H1 variants implies that this mode confers a selective advantage. More broadly, it suggests that distinct H1 isoforms can produce structurally non-equivalent 30-nm fibers, providing a framework for understanding how linker histone family expansion across eukaryotes is translated into chromatin architectural diversity.

Our structures reveal a hierarchy of higher-order chromatin organization driven by molecularly distinct mechanisms at each level. Within individual fibers, H1.3-mediated lateral bridging establishes the fundamental dinucleosomal stacking unit, producing the two-start helical fiber architecture resolved here. Between fibers, a distinct mechanism driven by H3.3 and H2A.W mediates back-to-back fiber dimerization. These two levels of organization are mechanistically separable: H1.3 governs intra-fiber nucleosomal stacking, whereas H3.3 N-terminal tails and H2A.W C-terminal extensions appear to govern inter-fiber contacts. The H3.3-specific A32T substitution at the dimerization interface, absent in canonical H3.1, likely contributes to stabilizing inter-fiber interactions, though the individual contribution of each variant remains to be resolved. This structural hierarchy, in which dinucleosomal stacking units are further organized into higher-order fiber assemblies, represents the type of multivalent interaction architecture proposed to underlie chromatin condensate formation (*18*). The lateral H1.3 bridging mode and the H3.3-H2A.W-dependent dimerization thus provide candidate structural mechanisms for generating the inter-fiber contacts that drive condensate assembly. The lateral bridging mode may further contribute to the reported role of H1.3 in DNA methylation partitioning (*24*), potentially by establishing compacted dinucleosomal units that restrict access of DNA methylation maintenance machsinery in a manner coupled to H1.3 occupancy.

Genome-wide profiling of Arabidopsis H1 variants indicates that H1.3 occupies both euchromatic and heterochromatic regions (*21, 22*), consistent with its function as a condition-dependent chromatin organizer. In euchromatic regions, stress-induced accumulation of H1.3 may promote localized dinucleosome stacking through the lateral bridging mode described here (Fig. 6A). We speculate that such stacking could restrict transcriptional machinery access by physically occluding promoter-proximal nucleosomes, providing a structural basis for H1.3-dependent gene repression under abiotic stress (*22, 23*). Nucleosome repeat length (NRL) is likely a key permissive factor governing this process. The lateral bridging mode is observed in fibers assembled with the 12 × 177 bp DNA, which provide sufficient linker DNA for the H1.3_B GD to reach the neighboring nucleosome. Shorter nucleosome repeats, characteristic of actively transcribed regions, may spatially confine H1.3 to the canonical on-dyad configuration and preclude lateral bridging (*14*). This NRL-dependent gating provides a framework through which the same H1.3 molecule produces structurally distinct outcomes depending on local chromatin context. In heterochromatin, H1.3 likely operates in concert with H2A.W and H3.3 to drive higher-order compaction through the fiber dimerization mechanism described above (Fig. 6B), providing a plausible structural basis for the formation of densely packed heterochromatic chromocenters essential for robust silencing of transposable elements.

**Fig. 6.**
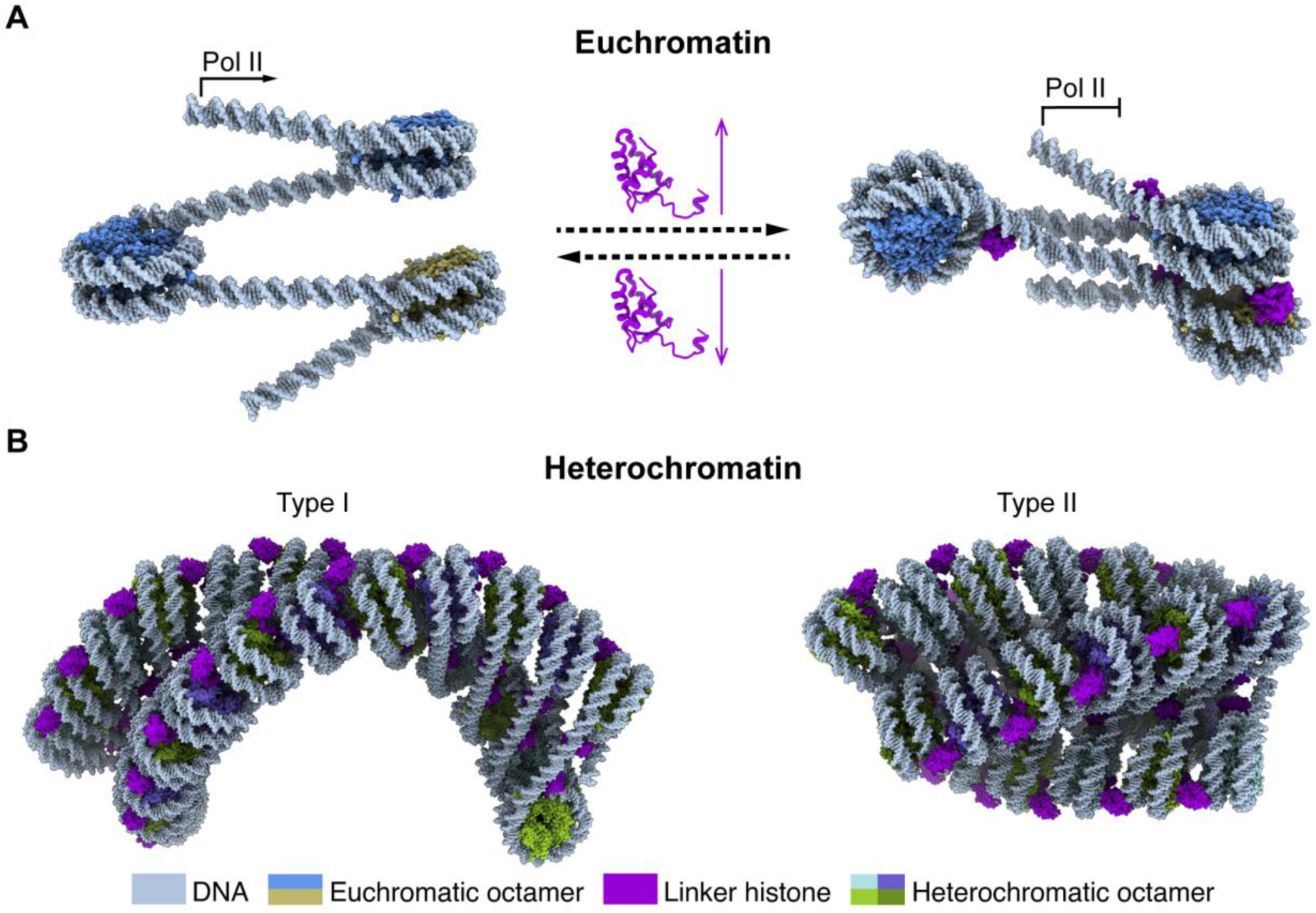
Proposed model for linker histone-mediated regulation of chromatin architecture within euchromatin and heterochromatin. (**A**) Model A illustrating the potential role of H1.3 in euchromatic regions. In actively transcribed loci (left), RNA polymerase II (Pol II) can traverse nucleosomes during transcription elongation. Upon stress-induced accumulation of H1.3 (right), the linker histone may dynamically bridge the adjacent nucleosomes to form dinucleosome units, which may create a structural barrier that limits access of Pol II or transcriptional activators, thereby modulating gene transcription. (**B**) Model B showing linker histones-mediated higher-order chromatin organizations in heterochromatin. H1.3 supports the formation of zig-zag 30-nm chromatin fiber through dinucleosome stacking (Type I). In the presence of histone variants such as H2A.W and H3.3, chromatin fibers may further undergo back-to-back dimerization to generate higher-order assemblies (Type II), potentially contributing to the formation of compacted heterochromatin.

Together, our results uncover an unanticipated mode of linker histone engagement in which H1.3 acts as a dynamic and multivalent scaffold, linking nucleosomes within and between chromatin fibers. This mechanism expands our understanding of how linker histones contribute to higher-order chromatin organization and highlights their adaptable roles in shaping chromatin architecture *in vivo*.

## Acknowledgments

We thank Dr. Guohong Li for helpful discussion and providing the plasmid containing 12 repeats of the 177-bp Widom 601 DNA sequence and. We thank Dr. Benjamin D. Engel for insightful discussion and valuable advice on cryo-electron tomography sample preparation and data analysis. We thank the staff members of the State Key Laboratory of Genetics and Development of Complex Phenotypes, School of Life Sciences, Fudan University, and the Cryo-EM System at the National Facility for Protein Science in Shanghai (NFPS), Shanghai Advanced Research Institute, Chinese Academy of Sciences, for providing technical support and assistance in data collection. We acknowledge the use of artificial intelligence-assisted tools for language editing of the manuscript. All scientific interpretations, data analyses, and conclusions were determined by the authors.

## Author contributions

Conceptualization: Y.W. and A.D.; Methodology: Y.W. and H.Z.; Investigation: Y.W., S.Y., J.W., X.W., D.Z., P.L., R.Z., J.S., H.Z. and J.G.; Visualization: Y.W.; Funding acquisition: A.D.; Project administration: A.D.; Supervision: J.Z., J.L., H.Z., J.G. and A.D.; Writing - original draft: Y.W.; Writing - review & editing: Y.W., S.Y., H.Z., J.G. and A.D.

## Data and materials availability

The electron microscopy raw data, density maps and atomic coordinates generated in this study have been deposited in the Electron Microscopy Public Image Archive (EMPIAR), the Electron Microscopy Data Bank (EMDB) and the Protein Data Bank (PDB), respectively. Accession numbers: 152_H2A-H3.3 (raw data: EMPIAR-12571, C1 map: EMD-62043, C2 map: EMD-62044, PDB: 9K44), 152_H2A.Z-H3.3 (raw data: EMPIAR-12572, C1 map: EMD-62045, C2 map: EMD-62046, PDB: 9K45), 152_H2A.W-H3.3 (raw data: EMPIAR-12573, C1 map: EMD-62047, C2 map: EMD-62048, PDB: 9K46), 177_12_H2A.Z (raw data: EMPIAR-13419, maps: EMD-67700, EMD-67701, EMD-67702, EMD-67703, EMD-67704, EMD-67705, EMD-67706; PDB: 21IU, 21IV), 177_12_H2A (maps: EMD-67707, EMD-67708, EMD-67709, EMD-67710, EMD-67711; PDB: 21IW), 177_12_H2A.W (maps: EMD-67712, EMD-67713, EMD-67714, EMD-67715, EMD-67716; PDB: 21IX), 177_12_H2A.Z_H5 (maps: EMD-67717, EMD-67718, EMD-67719, EMD-67720; PDB: 21IY, 21IZ), 177_12_H2A.W_H3.3 (maps: EMD-67721, EMD-67722, EMD-67723, EMD-67724, EMD-67725, EMD-67726, EMD-67727, EMD-67728, EMD-68924; PDB: 21JA, 21JB), and cryo-ET data (tomogram: EMD-67852).

## Supplementary Materials

Materials and Methods

Figures S1 to S15

Tables S1 to S9

