## Supplemental Figures 1-15 and supplemental tables 1-9 for "A lateral linker histone binding mode scaffolds dinucleosome stacking in chromatin fibers"

**This PDF file includes:**

Materials and Methods

Supplementary Figures S1-S15

Tables S1-S9

### Materials and methods

#### Histone purification, DNA preparation, and chromatin fiber assembly

The *Arabidopsis thaliana* H2A.Z (UniProt: Q9C944), H2A (Q9LD28), H2A.W (Q9FJE8), H2B (Q9LQQ4), H3.1 (P59226), H3.3 (P59169), H4 (P59259), linker histone H1.3 (P94109), and *Gallus gallus* linker histone H5 (P02259) were used for chromatin fiber reconstitution. Purification of histone dimers and tetramers, assembly of histone octamers, and nucleosome reconstitution were performed following the previously established protocols (*11*). Briefly, Arabidopsis H2A.Z-H2B, H2A-H2B and H2A.W-H2B dimers were separately co-expressed in *Escherichia coli*, as were H3.1-H4 and H3.3-H4 tetramers. Histone dimers and tetramers were purified by cation-exchange chromatography. Purified H3-H4 tetramers and H2A-H2B dimers were mixed at a molar ratio of 1:2 and incubated on ice, followed by size-exclusion chromatography to isolate correctly assembled histone octamers. Linker histones Arabidopsis H1.3 or *Gallus gallus* H5 were expressed in *E. coli* with an N-terminal SUMO tag and a C-terminal hexahistidine tag for purification. Linker histones were purified by nickel-affinity chromatography.

Preparation of the 12-repeat 177-bp Widom 601 DNA and assembly of chromatin fibers containing linker histones followed the protocol established by Dr. Guohong Li's laboratory (*13*). The full nucleotide sequence of the 12-repeat 177-bp Widom 601 DNA fragment used for chromatin fiber assembly in this study is provided below. Within each 177-bp repeat, the 147-bp Widom 601 core nucleosome-positioning sequence is underlined. The 177-bp×12 DNA fragment was obtained by restriction enzyme digestion of the corresponding plasmid. Chromatin fibers were assembled by salt gradient dialysis, gradually reducing ionic strength from 2 M to 0 M NaCl. Linker histones were added at a salt concentration of 0.6 M NaCl during dialysis.

ATCACGCGGCCGCCCTGGAGAATCCCGGTGCCGAGGCCGCTCAATTGGTCGTAGAC
AGCTCTAGCACCGCTTAAACGCACGTACGCGCTGTCCCCCGCGTTTTAACCGCCAAG
GGGATTACTCCCTAGTCTCCAGGCACGTGTCAGATATATACATCCTGTGCATGTAAG
ATCCAGTACTACGCGGCCGCCCTGGAGAATCCCGGTGCCGAGGCCGCTCAATTGGTC
GTAGACAGCTCTAGCACCGCTTAAACGCACGTACGCGCTGTCCCCCGCGTTTTAACC
GCCAAGGGGATTACTCCCTAGTCTCCAGGCACGTGTCAGATATATACATCCTGTGCA
TGTAAGATCCAGTACTACGCGGCCGCCCTGGAGAATCCCGGTGCCGAGGCCGCTCA
ATTGGTCGTAGACAGCTCTAGCACCGCTTAAACGCACGTACGCGCTGTCCCCCGCGT
TTTAACCGCCAAGGGGATTACTCCCTAGTCTCCAGGCACGTGTCAGATATATACATC
CTGTGCATGTAAGATCCAGTACTACGCGGCCGCCCTGGAGAATCCCGGTGCCGAGG
CCGCTCAATTGGTCGTAGACAGCTCTAGCACCGCTTAAACGCACGTACGCGCTGTCC
CCCGCGTTTTAACCGCCAAGGGGATTACTCCCTAGTCTCCAGGCACGTGTCAGATAT
ATACATCCTGTGCATGTAAGATCCAGTACTACGCGGCCGCCCTGGAGAATCCCGGTG
CCGAGGCCGCTCAATTGGTCGTAGACAGCTCTAGCACCGCTTAAACGCACGTACGC
GCTGTCCCCCGCGTTTTAACCGCCAAGGGGATTACTCCCTAGTCTCCAGGCACGTGT
CAGATATATACATCCTGTGCATGTAAGATCCAGTACTACGCGGCCGCCCTGGAGAAT
CCCGGTGCCGAGGCCGCTCAATTGGTCGTAGACAGCTCTAGCACCGCTTAAACGCAC
GTACGCGCTGTCCCCCGCGTTTTAACCGCCAAGGGGATTACTCCCTAGTCTCCAGGC
ACGTGTCAGATATATACATCCTGTGCATGTAAGATCCAGTACTACGCGGCCGCCCTG
GAGAATCCCGGTGCCGAGGCCGCTCAATTGGTCGTAGACAGCTCTAGCACCGCTTA
AACGCACGTACGCGCTGTCCCCCGCGTTTTAACCGCCAAGGGGATTACTCCCTAGTC
TCCAGGCACGTGTCAGATATATACATCCTGTGCATGTAAGATCCAGTACTACGCGGC
CGCCCTGGAGAATCCCGGTGCCGAGGCCGCTCAATTGGTCGTAGACAGCTCTAGCA

CCGCTTAAACGCACGTACGCGCTGTCCCCCGCGTTTTAACCGCCAAGGGGATTACTC
CCTAGTCTCCAGGCACGTGTCAGATATATACATCCTGTGCATGTAAGATCCAGTACT
ACGCGGCCGCCCTGGAGAATCCCGGTGCCGAGGCCGCTCAATTGGTCGTAGACAGC
TCTAGCACCGCTTAAACGCACGTACGCGCTGTCCCCCGCGTTTTAACCGCCAAGGGG
ATTACTCCCTAGTCTCCAGGCACGTGTCAGATATATACATCCTGTGCATGTAAGATC
CAGTACTACGCGGCCGCCCTGGAGAATCCCGGTGCCGAGGCCGCTCAATTGGTCGT
AGACAGCTCTAGCACCGCTTAAACGCACGTACGCGCTGTCCCCCGCGTTTTAACCGC
CAAGGGGATTACTCCCTAGTCTCCAGGCACGTGTCAGATATATACATCCTGTGCATG
TAAGATCCAGTACTACGCGGCCGCCCTGGAGAATCCCGGTGCCGAGGCCGCTCAAT
TGGTCGTAGACAGCTCTAGCACCGCTTAAACGCACGTACGCGCTGTCCCCCGCGTTT
TAACCGCCAAGGGGATTACTCCCTAGTCTCCAGGCACGTGTCAGATATATACATCCT
GTGCATGTAAGATCCAGTACTACGCGGCCGCCCTGGAGAATCCCGGTGCCGAGGCC
GCTCAATTGGTCGTAGACAGCTCTAGCACCGCTTAAACGCACGTACGCGCTGTCCCC
CGCGTTTTAACCGCCAAGGGGATTACTCCCTAGTCTCCAGGCACGTGTCAGATATAT
ACATCCTGTGCATGTAAGATCTGAT

**Cryo-EM grid preparation and single-particle data collection**

Assembled chromatin fibers were cross-linked on ice using 0.5% glutaraldehyde for 15 min to stabilize chromatin fibers during vitrification. Sample concentration was assessed by negative-stain transmission electron microscopy at room temperature. Cross-linked samples at suitable concentrations were vitrified using a Vitrobot Mark IV (Thermo Fisher Scientific) by rapid plunging into liquid ethane. Graphene oxide-coated grids with R1.2/1.3 geometry were used without glow discharge. Cryo-EM data were collected on a Titan Krios G4 microscope (Thermo

Fisher Scientific) equipped with a Falcon 4i direct electron detector and a Selectris X energy filter, operated at 300 keV. Data were acquired using Smart EPU software (Thermo Fisher Scientific) in EER format at a nominal magnification of 130,000×, corresponding to a calibrated pixel size of 0.959 Å. The defocus range was set from -0.6 to -1.6 µm, with a total electron dose of 40 $e^{-}/Å^2$.

**Cryo-EM single-particle data processing**

Cryo-EM data were processed using RELION v4.0 (*44*) and cryoSPARC v4.7 (*45*). In RELION, movie frames were motion-corrected using RELION's implementation of motion correction. Micrographs were filtered based on defocus values, estimated resolution, and astigmatism. Reference-free particle picking was performed, followed by multiple rounds (typically three) of 2D classification. Selected particles were subjected to iterative 3D classification. The initial 3D classification used EMD-2600 (*13*) as a reference, whereas subsequent rounds used maps reconstructed in this study. After each round of 3D classification, selected particles were subjected to additional 2D classification before proceeding to the next round. Final particles were refined by 3D auto-refinement and post-processing. For the 177_12_H2A.Z dataset, maps showing all 12 nucleosomes were reconstructed in RELION. For the back-to-back fiber dimerization observed in the 177_12_H2A.W_H3.3 dataset, particles were exported from RELION and further processed in cryoSPARC. In cryoSPARC, movies were motion-corrected using Patch Motion Correction. Micrographs were filtered using criteria similar as those applied in RELION. Template-based particle picking was performed using 2D templates generated from RELION reconstructions. After multiple rounds of 2D classification, particles were subjected to ab-initio reconstruction and heterogeneous refinement. Selected classes were further refined using homogeneous refinement and non-uniform refinement, followed by local refinement with appropriate masks. Because linker histones lack intrinsic symmetry, whereas nucleosomes exhibit pseudo-C2 symmetry, symmetry-breaking features were observed, including apparent binding of two linker histone molecules at a single nucleosome. For H5-containing chromatin fibers, symmetry expansion (C2) was applied prior to 3D classification. For other datasets, this issue was mitigated by 3D classification.

**Model building and visualization**

The nucleosome atomic model (PDB ID: 9K3Z) was used as the initial model. Atomic models of linker histones H1.3 and H5 were predicted using AlphaFold 3 (*46*) and used as starting models. Models were docked into cryo-EM maps using UCSF Chimera (*47*). Real-space refinement was performed using PHENIX (*48*), and manual model adjustment was carried out in Coot (*49*). For the 177_12_H2A.Z dataset, a composite map was generated by combining two focused mononucleosome maps using the PHENIX composite map procedure (*50*). Cryo-EM density maps were visualized using UCSF ChimeraX (*51*). Atomic models were rendered using Open-Source-PyMOL (*52*) and ChimeraX.

**Quantitative analysis of chromatin fiber geometry**

Quantitative measurements of chromatin fiber geometry were performed using ChimeraX. For each dataset, nucleosome atomic models were first docked into the corresponding chromatin fiber cryo-EM density maps. In the Arabidopsis H2A.Z-H3.1-containing chromatin fiber, all twelve nucleosomes were clearly resolved and included in the analysis. In the H2A.W-H3.3-containing chromatin fiber, ten nucleosomes were visible, but two nucleosomes with poorly resolved density were excluded, the remaining eight nucleosomes were used for quantitative measurements. For the H2A.W-H3.3-containing back-to-back fiber dimerization, eight nucleosomes from each individual fiber were included in the analysis. Geometric parameters were defined based on the centers and orientations of histone octamers and tetramers. The centers of histone octamers and tetramers, the tetramer axis, and the dinucleosome axis were defined in ChimeraX. The dinucleosome axis was defined using the tetramer axes of two adjacent nucleosomes. Inter-nucleosome distances, including nucleosome stacking distance and dinucleosome distance, were calculated as the distance between the centers of adjacent histone octamers. Nucleosome stacking angles were calculated using the axes of the tetramers from two stacked nucleosomes. Nucleosome stacking rotation angles were calculated based on four reference points: the centers of the two histone octamers and the centers of the two corresponding tetramers. For adjacent dinucleosome units, the opening angle was calculated using the tetramer axes of the two dinucleosomes. The linker twist angle between adjacent dinucleosomes was calculated using four reference points defined by the octamer and tetramer centers. The rotation angle between neighboring dinucleosome units was calculated using the axes defined by the tetramers of one dinucleosome and those of the adjacent dinucleosome.

**Rice protoplast preparation and cryo-ET sample vitrification**

Rice protoplasts were prepared following the standard enzymatic digestion protocol. Briefly, rice leaf tissues were cut into fine strips and incubated in enzyme solution containing cellulase and macerozyme to release protoplasts. Protoplasts were filtered, washed, and resuspended in isotonic buffer. Protoplast concentration was adjusted by light microscopy before vitrification. Protoplast samples were vitrified using an EM GP2 plunger (Leica Microsystems) by rapid freezing in liquid ethane. R1.2/1.3 200-mesh amorphous alloy film grids were used and glow-discharged in a Pelco easiGlow (Ted Pella) at 0.39 mbar and 15 mA for 30 s prior to sample application.

**Cryo-ET lamella preparation and data collection**

Vitrified grids were transferred to a cryogenic dual-beam focused ion beam/scanning electron microscope (Arctis, Thermo Fisher Scientific). Lamellae were generated by cryo-focused ion beam milling at a tilt angle of 15°. Grids containing lamellae were subsequently transferred to a 300-keV Titan Krios microscope for cryo-ET data collection using Tomography software (Thermo Fisher Scientific). Tilt series were acquired using a dose-symmetric scheme over an angular range of -70° to +40°, with 3° increments. Data were collected at a nominal magnification of 53,000×, corresponding to a pixel size of 2.4 Å, with a defocus range of -4 to -6 µm and an electron dose of 3 $e^-/Å^2$ per tilt.

**Cryo-ET data processing and visualization**

Tilt series were motion-corrected using Warp (*53*). Patch-based tilt-series alignment was performed via WarpTools (ts_etomo_patches), which calls IMOD/eTomo routines for patch tracking and alignment (*54*). Tomograms were reconstructed in Warp using weighted back projection with a final pixel size of 9.6 Å (bin 4). Denoising was performed using Warp Noise2Map. Visualization was carried out using IMOD (*54*) and the Artia plugin in ChimeraX. For illustrative purposes, a low-resolution mononucleosome density map was positioned at selected tomogram coordinates corresponding to nucleosome-like densities to visualize relative nucleosome stacking geometries in situ.

### Supplementary Figures

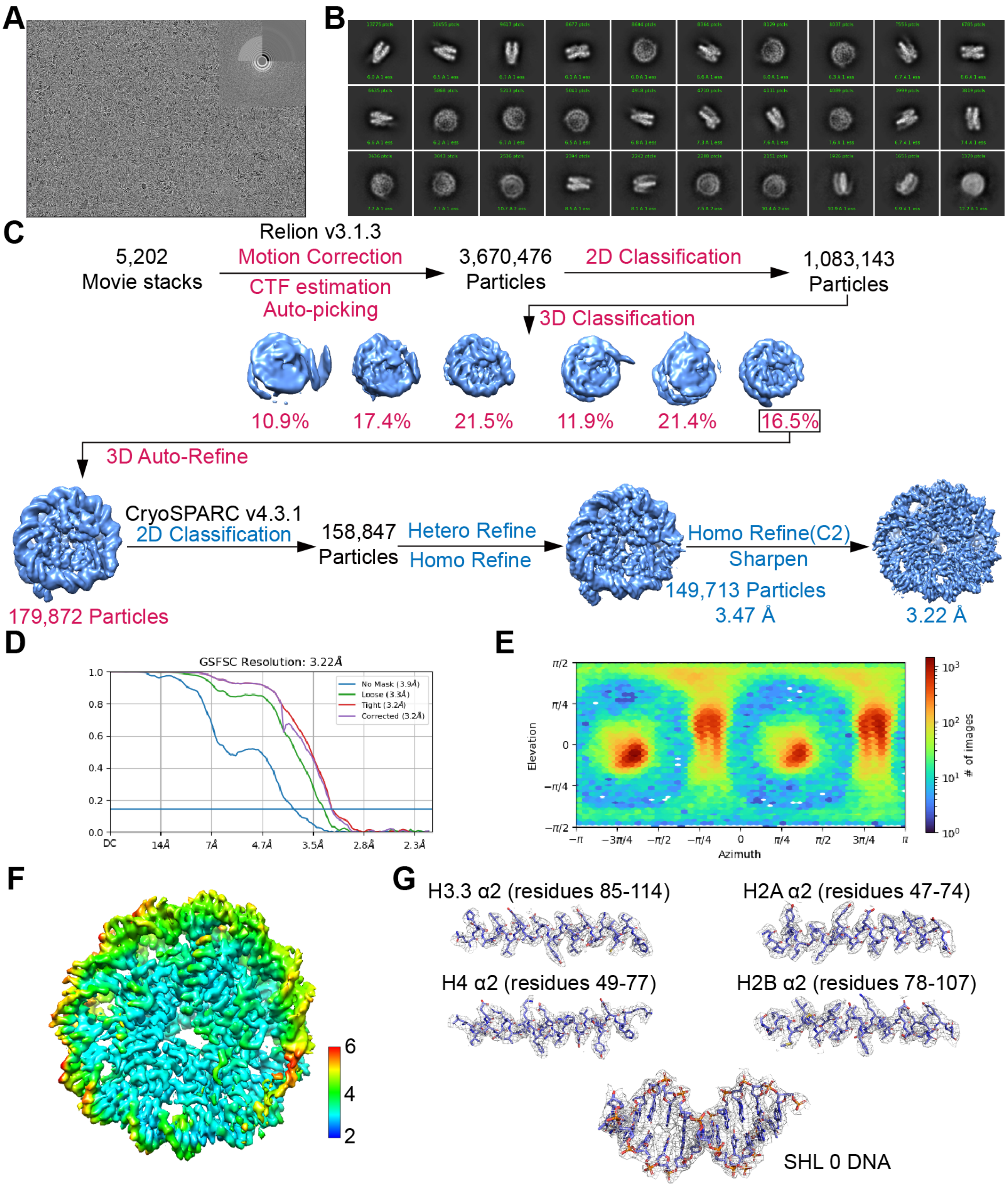


**Fig. S1. Cryo-EM data processing of nucleosomes containing Arabidopsis H2A, H2B, H3.3, H4 and native 152 DNA (152-H2A-H3.3)**

(**A**) Representative electron micrograph with inset Fourier transform (Upper-Right).

(**B**) Selected reference-free 2D class averages.

(**C**) Workflow of cryo-EM data processing of 152-H2A-H3.3.

(**D**) FSC curves of the C2 density map of 152-H2A-H3.3.

(**E**) Angular distribution of 152-H2A-H3.3 C2 density map.

(**F**) Local resolution estimation of 152-H2A-H3.3 C2 map.

(**G**) Map-to-model fits for 152-H2A-H3.3 representative regions: histone density from C2 map and DNA from C1 map.

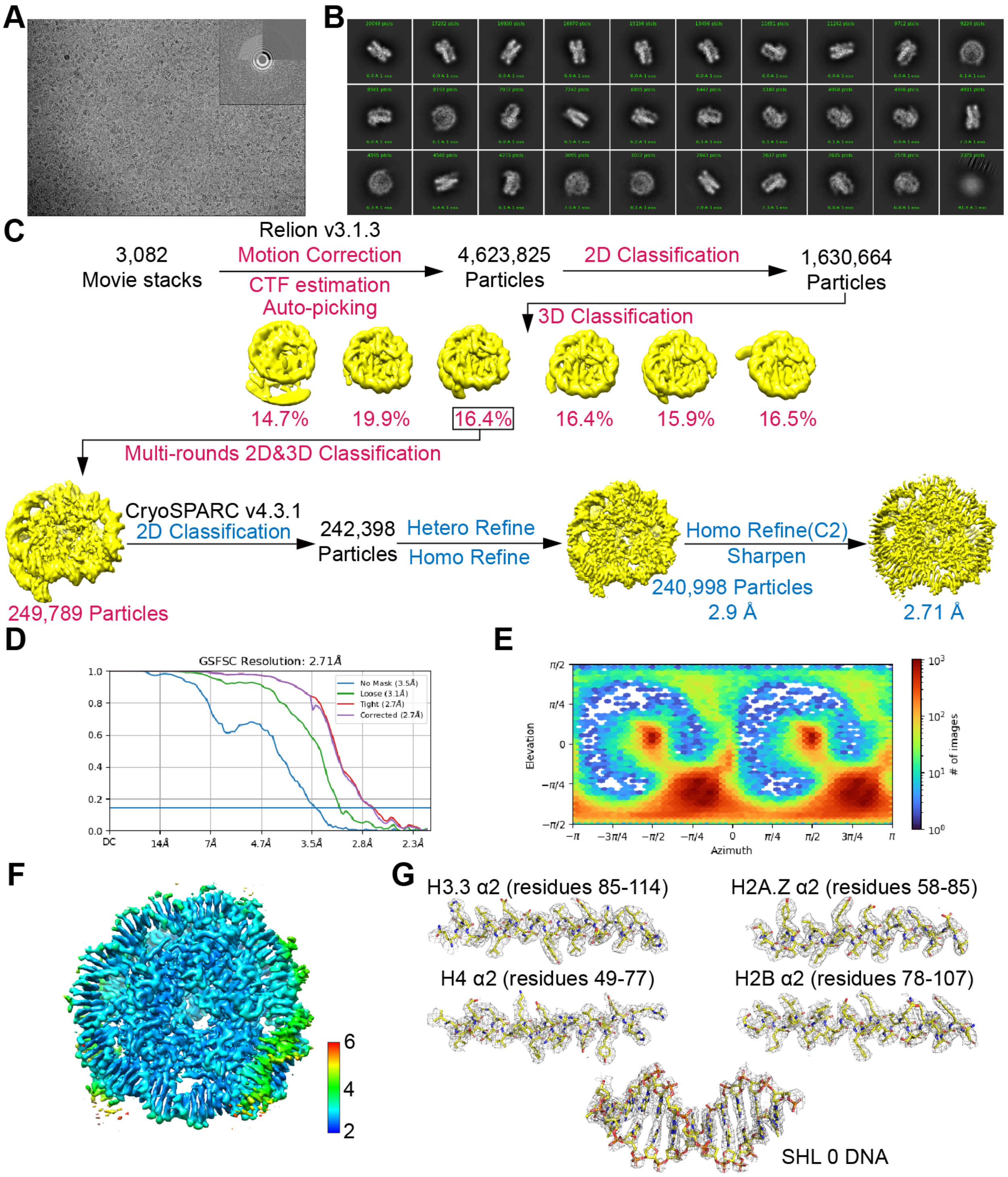


**Fig. S2. Cryo-EM data processing of nucleosomes containing Arabidopsis H2A.Z, H2B, H3.3, H4 and native 152 DNA (152-H2A.Z-H3.3)**

(**A**) Representative electron micrograph with inset Fourier transform (Upper-Right).

(**B**) Selected reference-free 2D class averages.

(**C**) Workflow of cryo-EM data processing of 152-H2A.Z-H3.3.

(**D**) FSC curves of the C2 density map of 152-H2A.Z-H3.3.

(**E**) Angular distribution of 152-H2A.Z-H3.3 C2 density map.

(**F**) Local resolution estimation of 152-H2A.Z-H3.3 C2 map.

(**G**) Map-to-model fits for 152-H2A.Z-H3.3 representative regions: histone density from C2 map and DNA from C1 map.

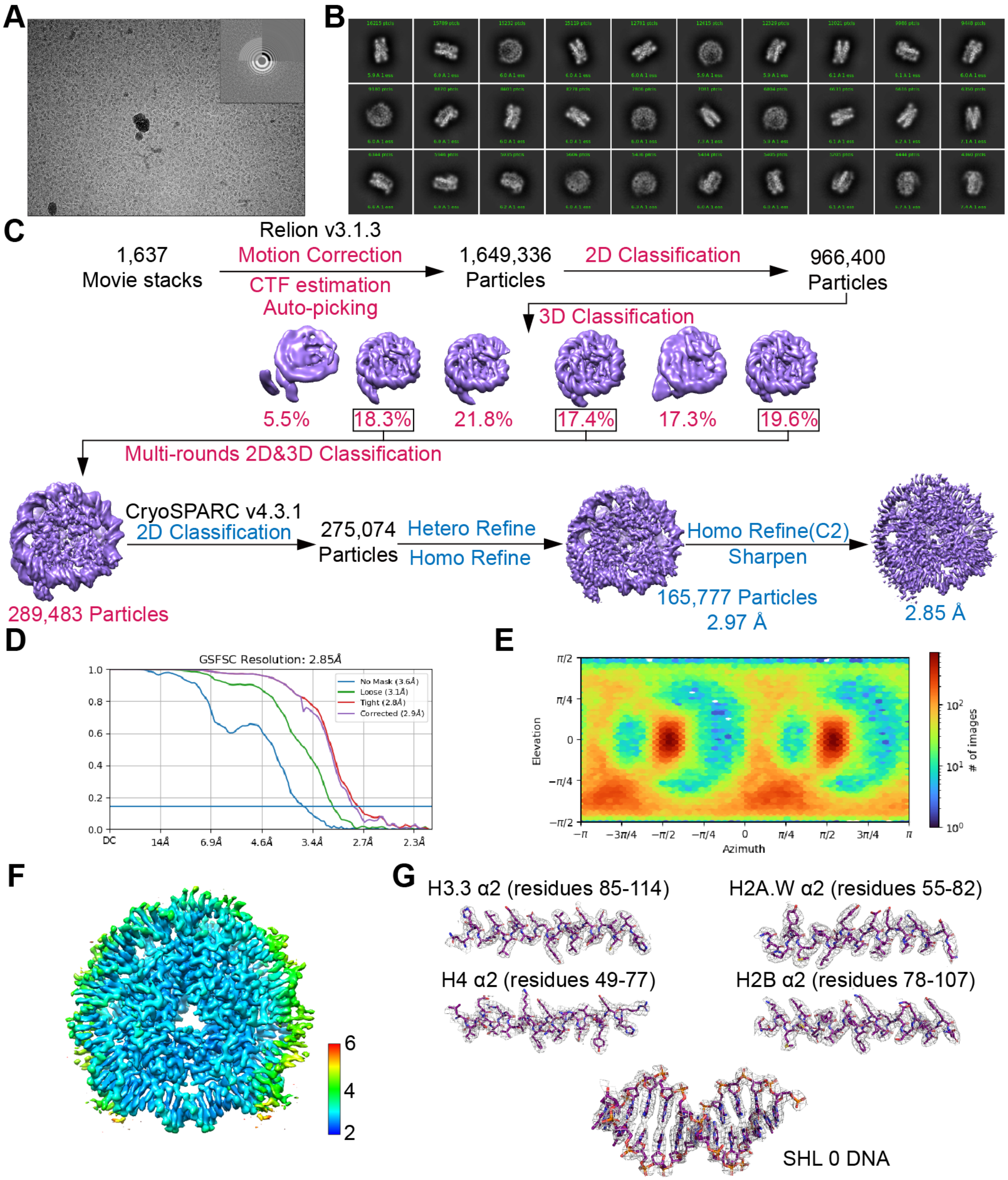


**Fig. S3. Cryo-EM data processing of nucleosomes containing Arabidopsis H2A.W, H2B, H3.3, H4 and native 152 DNA (152-H2A.W-H3.3)**

(**A**) Representative electron micrograph with inset Fourier transform (Upper-Right).

(**B**) Selected reference-free 2D class averages.

(**C**) Workflow of cryo-EM data processing of 152-H2A.W-H3.3.

(**D**) FSC curves of the C2 density map of 152-H2A.W-H3.3.

(**E**) Angular distribution of 152-H2A.W-H3.3 C2 density map.

(**F**) Local resolution estimation of 152-H2A.W-H3.3 C2 map.

(**G**) Map-to-model fits for 152-H2A.W-H3.3 representative regions: histone density from C2 map and DNA from C1 map.

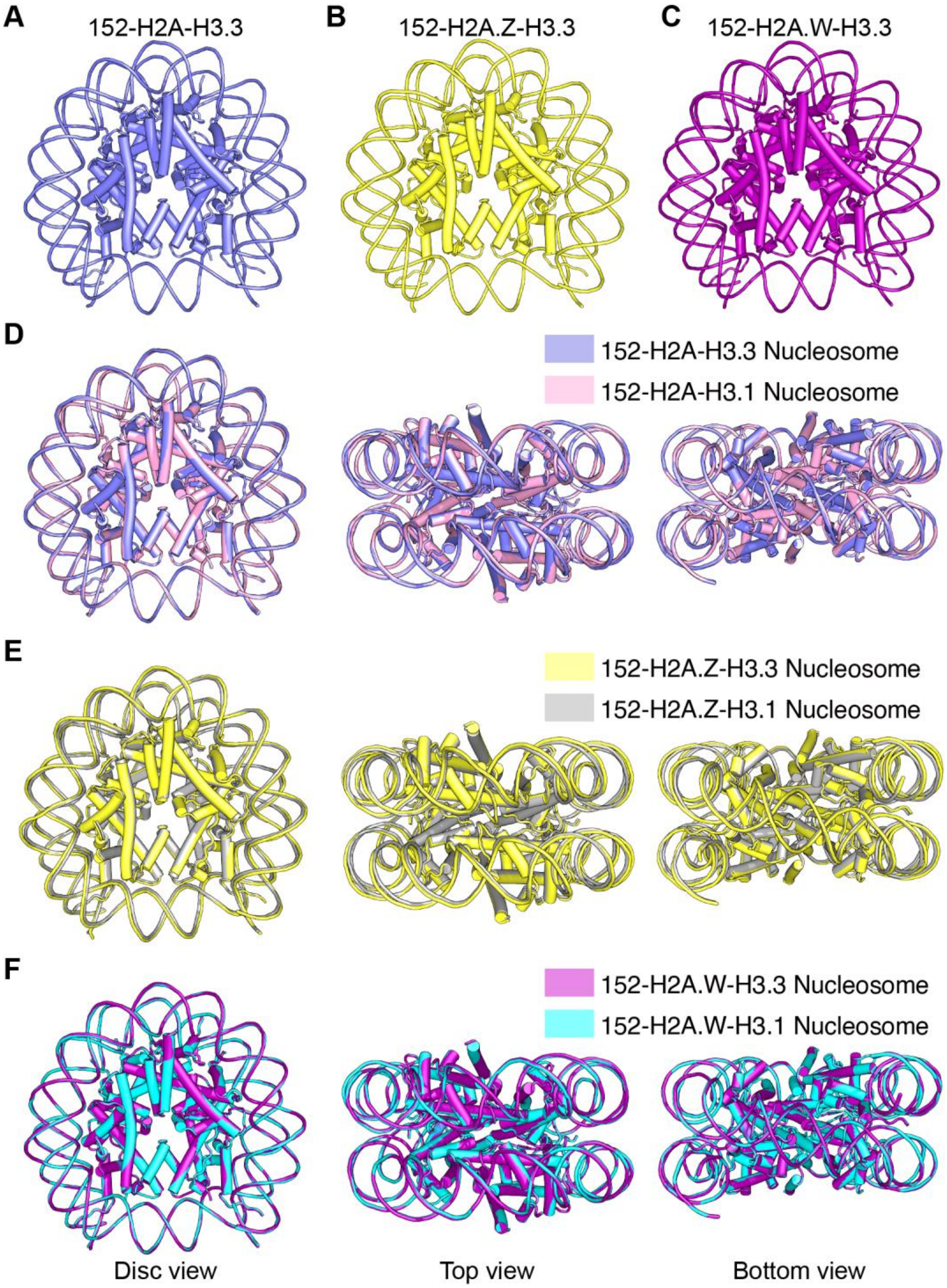


**Fig. S4. Comparison of Arabidopsis nucleosomes containing H2A/H2A.Z/H2A.W and H3.1/H3.3**

(**A-C**) Atomic models of the 152-H2A-H3.3 (A), 152-H2A.Z-H3.3 (B), and 152-H2A.W-H3.3 (C) nucleosomes shown in disc view. The 152 DNA is a Widom 601-like high-affinity nucleosome positioning sequence previously identified in Arabidopsis.

(**D-F**) Global structural superimpositions of nucleosomes containing H3.3 with their respective H3.1 counterparts. The overall architectures of the nucleosomes assembled with H3.1 or H3.3 are similar.

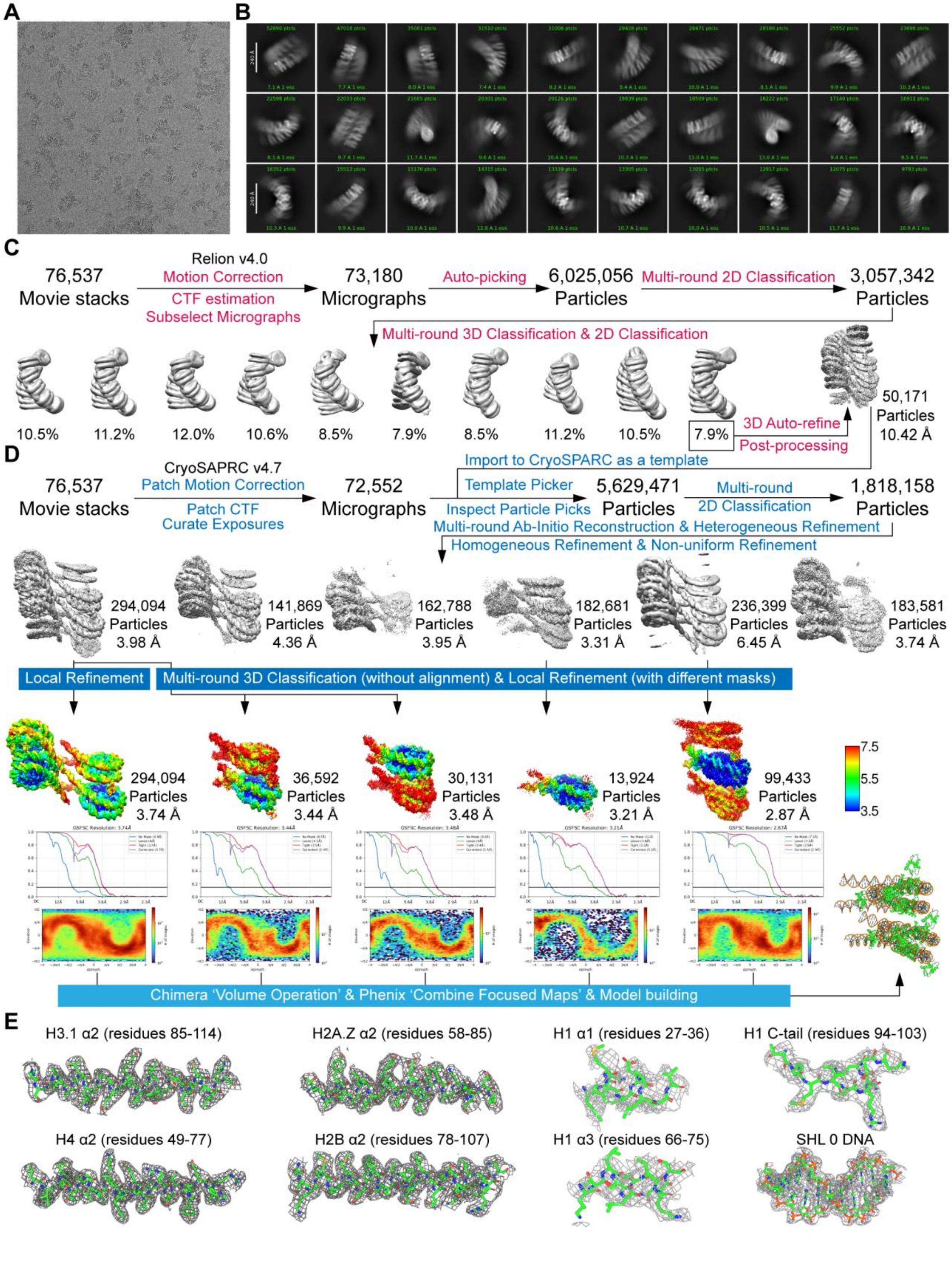
A
B
C
76,537 Movie stacks
Relion v4.0
Motion Correction
CTF estimation
Subselect Micrographs
73,180 Micrographs
Auto-picking
6,025,056 Particles
Multi-round 2D Classification
3,057,342 Particles
Multi-round 3D Classification & 2D Classification
10.5%
11.2%
12.0%
10.6%
8.5%
7.9%
8.5%
11.2%
10.5%
7.9%
3D Auto-refine
Post-processing
50,171 Particles 10.42 Å
D
76,537 Movie stacks
CryoSAPRC v4.7
Patch Motion Correction
Patch CTF
Curate Exposures
72,552 Micrographs
Import to CryoSPARC as a template
Template Picker
Inspect Particle Picks
5,629,471 Particles
Multi-round 2D Classification
1,818,158 Particles
Multi-round Ab-Initio Reconstruction & Heterogeneous Refinement
Homogeneous Refinement & Non-uniform Refinement
294,094 Particles 3.98 Å
141,869 Particles 4.36 Å
162,788 Particles 3.95 Å
182,681 Particles 3.31 Å
236,399 Particles 6.45 Å
183,581 Particles 3.74 Å
Local Refinement
Multi-round 3D Classification (without alignment) & Local Refinement (with different masks)
294,094 Particles 3.74 Å
36,592 Particles 3.44 Å
30,131 Particles 3.48 Å
13,924 Particles 3.21 Å
99,433 Particles 2.87 Å
7.5
5.5
3.5
Chimera 'Volume Operation' & Phenix 'Combine Focused Maps' & Model building
E
H3.1 α2 (residues 85-114)
H2A.Z α2 (residues 58-85)
H1 α1 (residues 27-36)
H1 C-tail (residues 94-103)
H4 α2 (residues 49-77)
H2B α2 (residues 78-107)
H1 α3 (residues 66-75)
SHL 0 DNA

**Fig. S5. Cryo-EM data processing of the chromatin fiber containing H1.3, H2A.Z, H2B, H3.1, H4 and 12-repeat 177-bp Widom 601 DNA (177_12_H2A.Z).**

(**A**) Representative cryo-EM micrograph collected using a Falcon 4i direct electron detector (4096 × 4096 pixels; pixel size, 0.959 Å).

(**B**) Selected reference-free two-dimensional (2D) class averages.

(**C**) Workflow of cryo-EM data processing performed in RELION v4.0.

(**D**) Workflow of cryo-EM data processing performed in cryoSPARC v4.7.

(**E**) Representative map-to-density fits for selected regions of the reconstructed chromatin fiber.

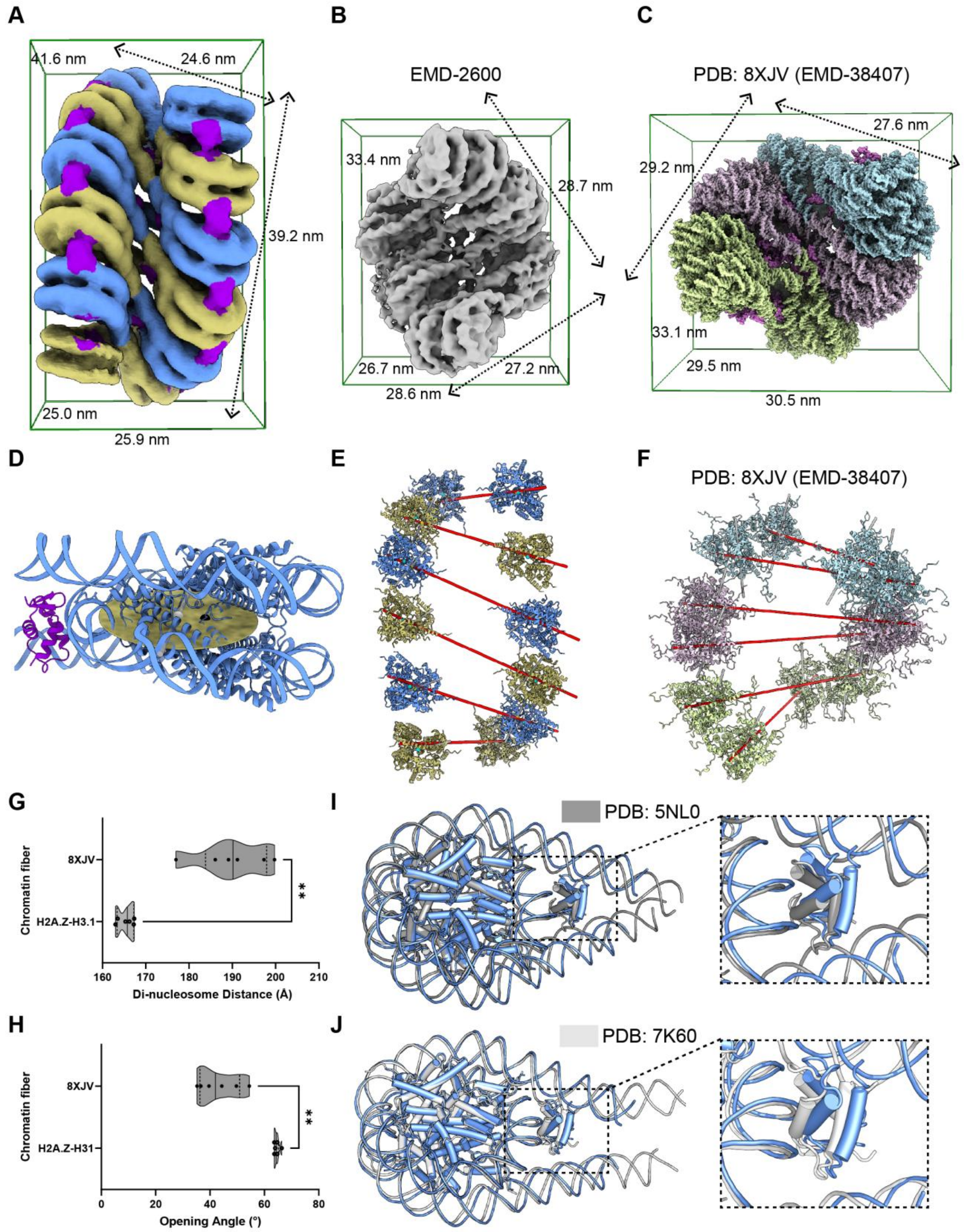


**Fig. S6. Structural comparison between the Arabidopsis 177_12_H2A.Z chromatin fiber and previously reported animal H5-chromatin fibers.**

(**A**) Dimensions of the cryo-EM density map of the Arabidopsis 177_12_H2A.Z chromatin fiber measured using the "Measure Volume and Area (Blob)" tool in UCSF ChimeraX. The map is rendered in ChimeraX and contoured at a level of 0.0035 (map units). Adjacent dinucleosome units are colored differently for clarity, with linker histone H1.3 highlighted in dark violet.
(**B**) Dimensions of the cryo-EM density map of EMD-2600, measured using the ChimeraX Blob tool. The map is rendered in ChimeraX and contoured at a level of 3.5 (map units).
(**C**) Dimensions of the atomic model of PDB 8XJV (corresponding map EMD-38407), measured using the ChimeraX Blob tool. Each tetranucleosomal unit is shown in a distinct color, with linker histone H5 highlighted in violet.
(**D**) Schematic definition of geometric parameters used for quantitative analysis of nucleosome stacking. The center of histone octamer (black dot), center of histone tetramer (white dot), tetramer axis and tetramer plane are defined and used to calculate inter-nucleosome distance, stacking angle, rotation angle, opening angle, and twist angle.
(**E, F**) Definitions of the mononucleosome axis and dinucleosome axis in the 177_12_H2A.Z chromatin fiber (E) and in PDB 8XJV (F). The dinucleosome axis was used to measure the inter dinucleosome angle between adjacent dinucleosome units. Red lines are axes of adjacent histone tetramers (nucleosomes N and N+1).
(**G, H**) Violin plots comparing distances (G) and opening angles (H) between adjacent nucleosomes (N and N+1) measured for the twelve nucleosomes in the Arabidopsis H2A.Z-H3.1-containing chromatin fiber and those measured for the chromatin fiber from PDB 8XJV. Statistical significance was assessed using the Mann–Whitney test ($P < 0.01$).
(**I, J**) Structural superposition of a single nucleosome unit from the Arabidopsis 177_12_H2A.Z chromatin fiber (PDB: 21IU) with the previously reported structures, PDB: 5NL0 (I) and PDB: 7K60 (J). The linker histone H1.3 bound at the nucleosomal on-dyad site exhibits a highly similar mode of DNA engagement.

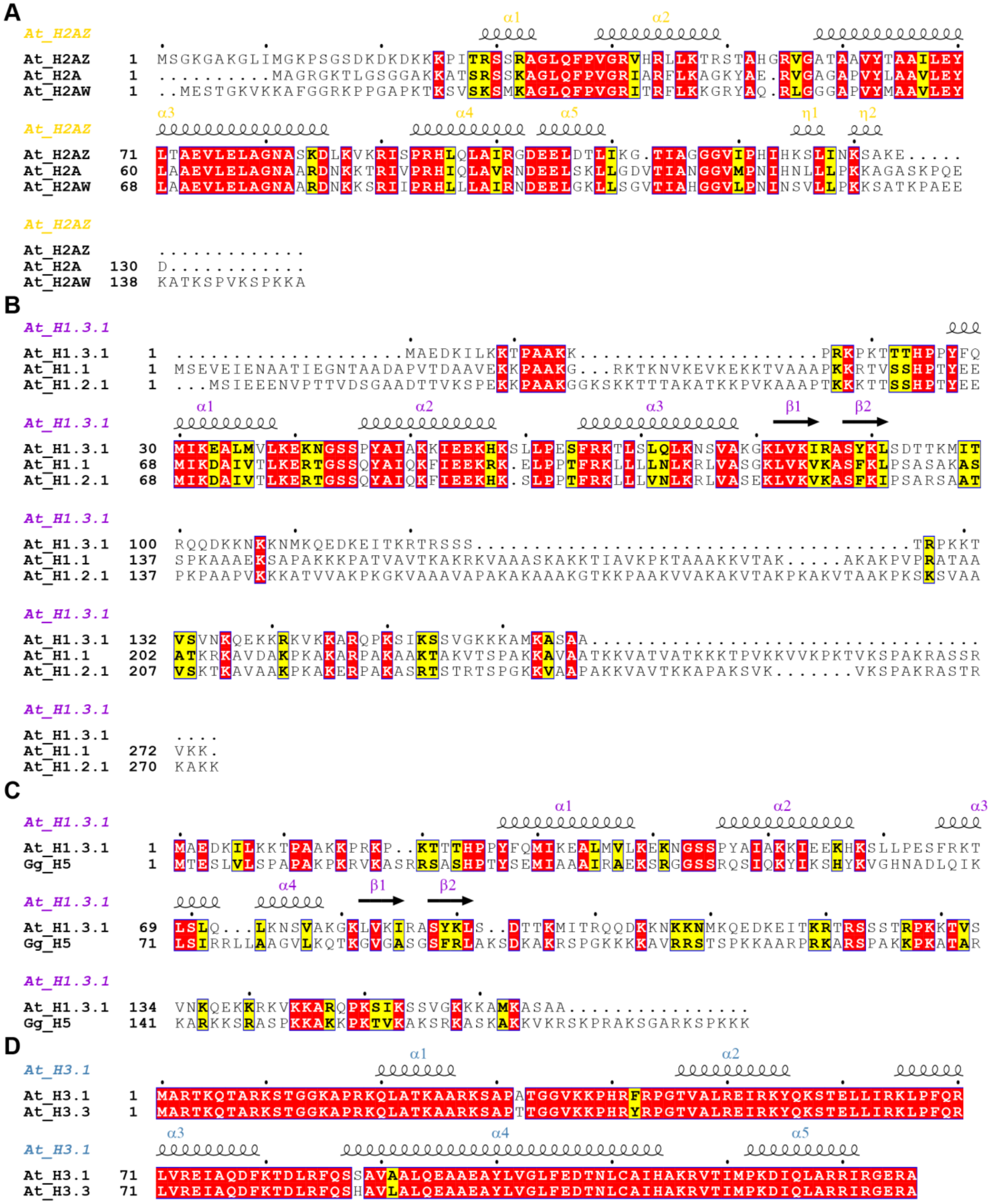


**Fig. S7. Sequence alignments of Arabidopsis histones used for chromatin fiber assembly in this study.**

(**A**) Sequence alignment of Arabidopsis H2A (Q9LD28), H2A.Z (Q9C944) and H2A.W (Q9FJE8).

(**B**) Sequence alignment of Arabidopsis linker histone H1.1 (A0A178W387), H1.2 (P26569) and H1.3 (P94109).

(**C**) Sequence alignment of Arabidopsis linker histone H1.3 (P94109) and *Gallus gallus* linker histone H5 (P02259).

(**D**) Sequence alignment of Arabidopsis H3.1 (P59226) and H3.3 (P59169).

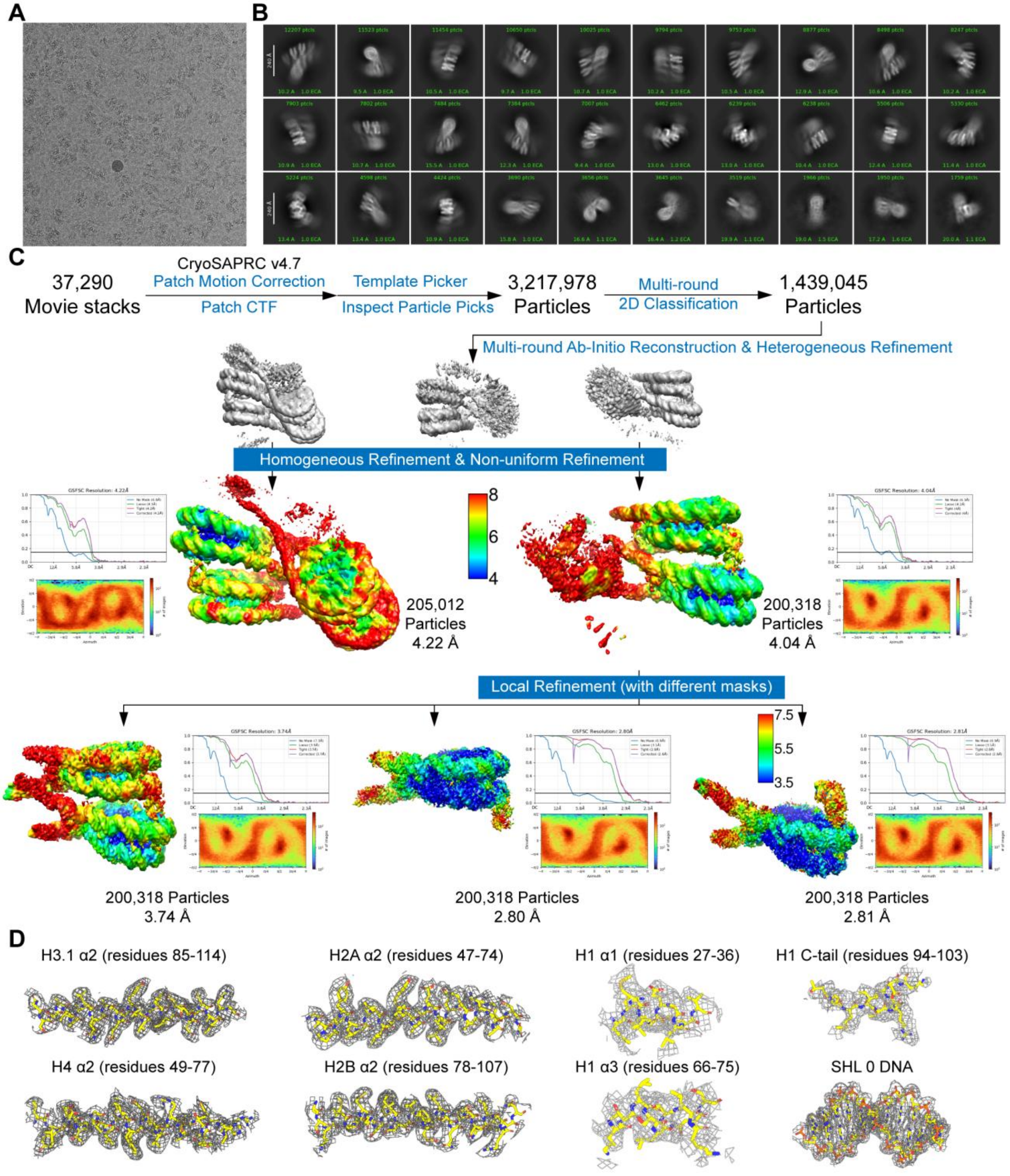


**Fig. S8. Cryo-EM data processing of the chromatin fiber containing H1.3, H2A, H2B, H3.1, H4 and 12-repeat 177-bp Widom 601 DNA (177_12_H2A).**

(**A**) Representative cryo-EM micrograph collected using a Falcon 4i direct electron detector (4096 × 4096 pixels; pixel size, 0.959 Å).

(**B**) Selected reference-free two-dimensional (2D) class averages.

(**C**) Workflow of cryo-EM data processing performed in cryoSPARC v4.7.

(**D**) Representative map-to-density fits for selected regions of the reconstructed chromatin fiber.

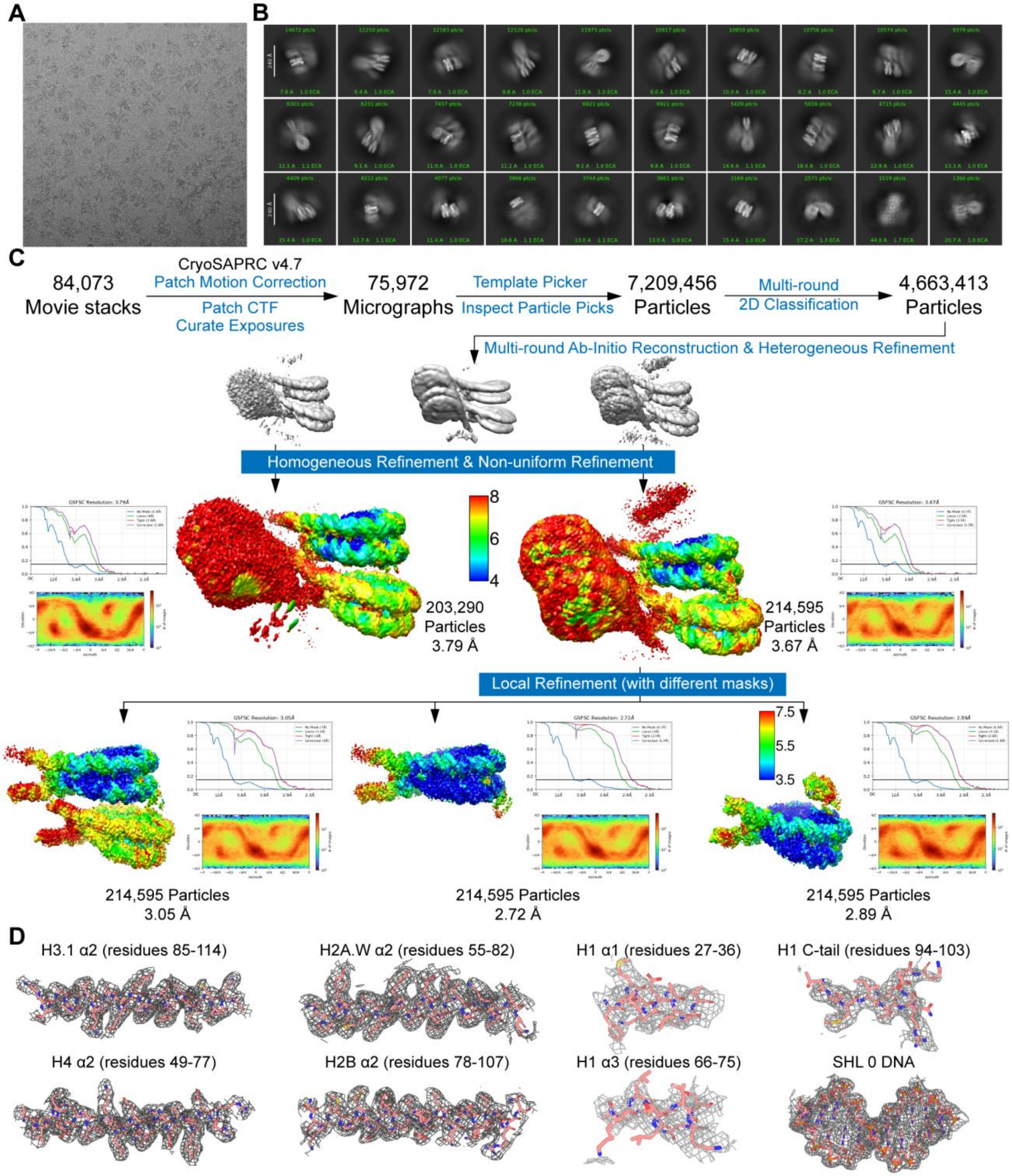


**Fig. S9. Cryo-EM data processing of the chromatin fiber containing H1.3, H2A.W, H2B, H3.1, H4 and 12-repeat 177-bp Widom 601 DNA (177_12_H2A.W).**

(**A**) Representative cryo-EM micrograph collected using a Falcon 4i direct electron detector (4096 × 4096 pixels; pixel size, 0.959 Å).

(**B**) Selected reference-free two-dimensional (2D) class averages.

(**C**) Workflow of cryo-EM data processing performed in cryoSPARC v4.7.

(**D**) Representative map-to-density fits for selected regions of the reconstructed chromatin fiber.

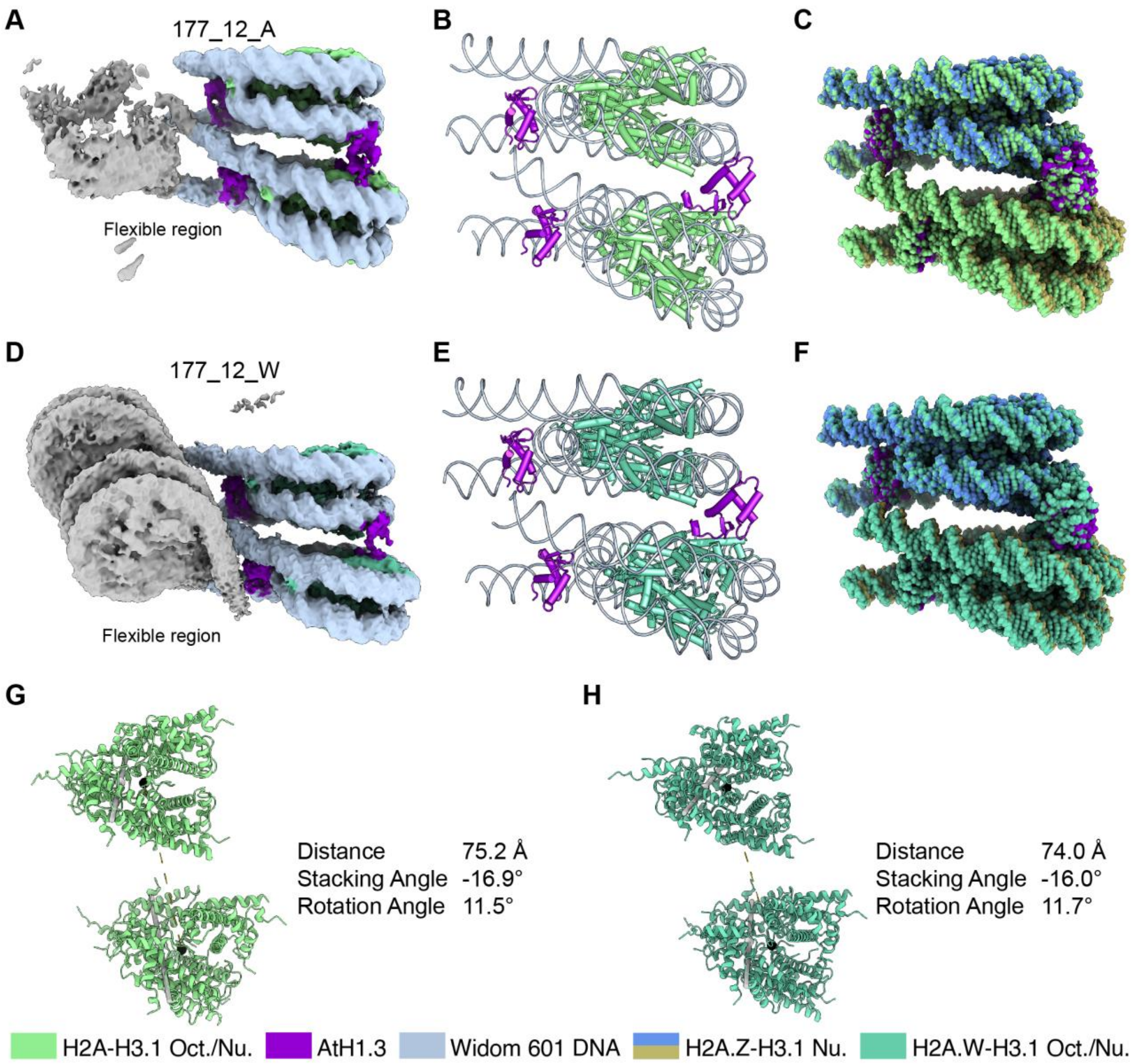


**Fig. S10. Structural basis of chromatin fibers containing H2A/H2A.W, H2B, H3.1, H4, and H1.3.**

**(A, D)** Cryo-EM density maps of chromatin fibers assembled with canonical H2A (A) or plant-specific variant H2A.W (D). Flexible regions are indicated. DNA, linker histone H1.3, and histone octamers are shown in distinct colors. The maps have been deposited in the Electron Microscopy Data Bank (EMDB) under accession numbers EMD-67711 (A) and EMD-67716 (D). Oct./Nu. indicates octamer/nucleosome.

**(B, E)** Atomic models of H2A- (B) or H2A.W-containing (E) dinucleosomes stabilized by linker histone H1.3. DNA, H1.3, and histone octamers are colored distinctly. For (B), the atomic model was built into the overall chromatin fiber map shown in (A) (EMD-67711), together with two focused maps deposited under accession numbers EMD-67709 and EMD-67710. For (E), the atomic model was built into the overall chromatin fiber map shown in (D) (EMD-67716), together with two focused maps deposited under accession numbers EMD-67714 and EMD-67715. The atomic coordinates have been deposited in the Protein Data Bank (PDB) under accession numbers 21IW (B) and 21IX (E).

**(C, F)** Structural alignment of the H2A-containing (C, light green) or H2A.W-containing (F, medium aquamarine) dinucleosome with the H2A.Z-containing dinucleosome (PDB: 21IV, cornflower blue, dark khaki and dark violet).

**(G, H)** Measurements of inter-nucleosomal distance, stacking angle, and rotation angle of H2A- (G) or H2A.W-containing (H) dinucleosome.

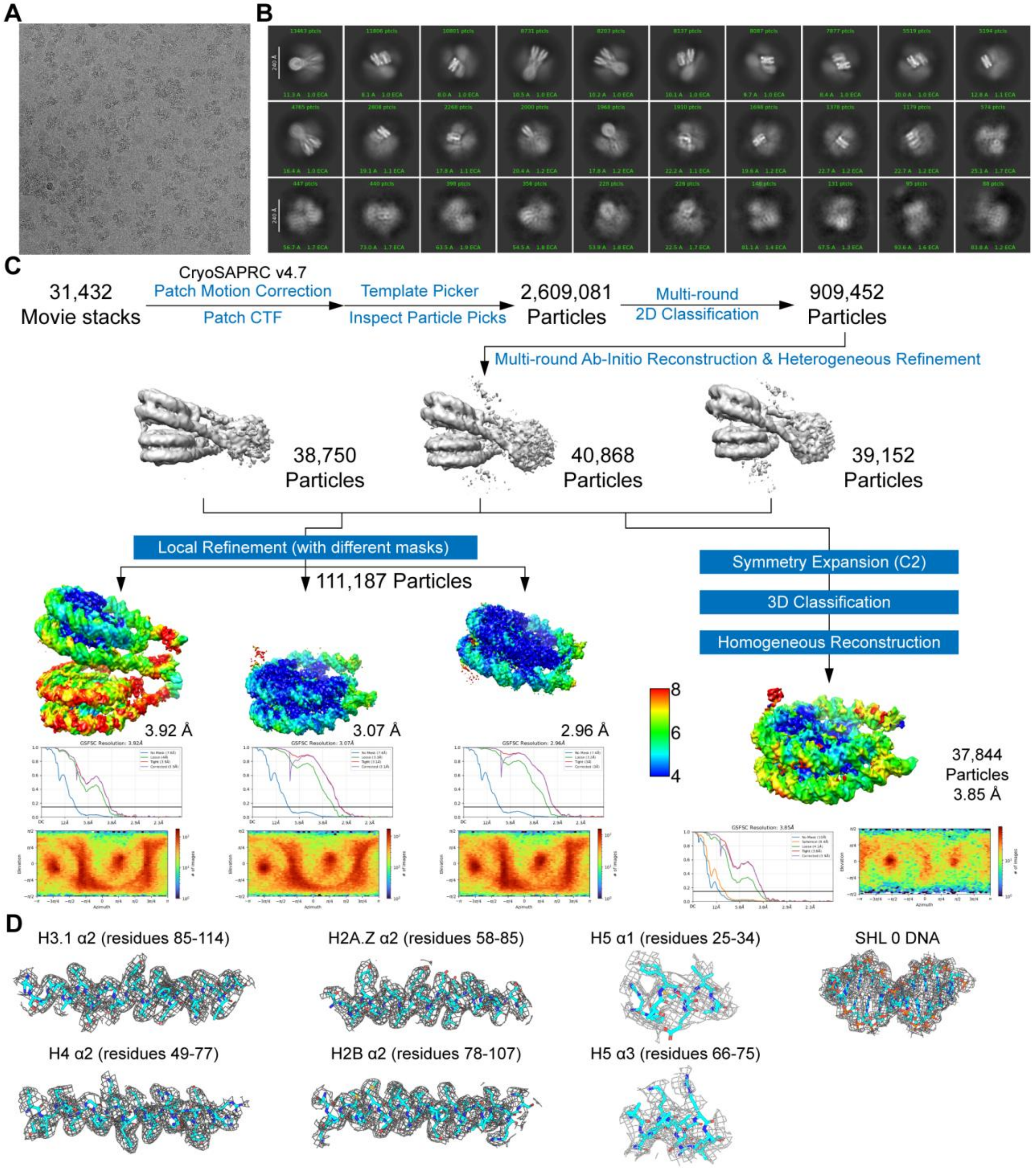


**Fig. S11. Cryo-EM data processing of the chromatin fiber containing H5, H2A.Z, H2B, H3.1, H4 and 12-repeat 177-bp Widom 601 DNA (177_12_H2A.Z_H5).**

(**A**) Representative cryo-EM micrograph collected using a Falcon 4i direct electron detector (4096 × 4096 pixels; pixel size, 0.959 Å).

(**B**) Selected reference-free two-dimensional (2D) class averages.

(**C**) Workflow of cryo-EM data processing performed in cryoSPARC v4.7.

(**D**) Representative map-to-density fits for selected regions of the reconstructed chromatin fiber.

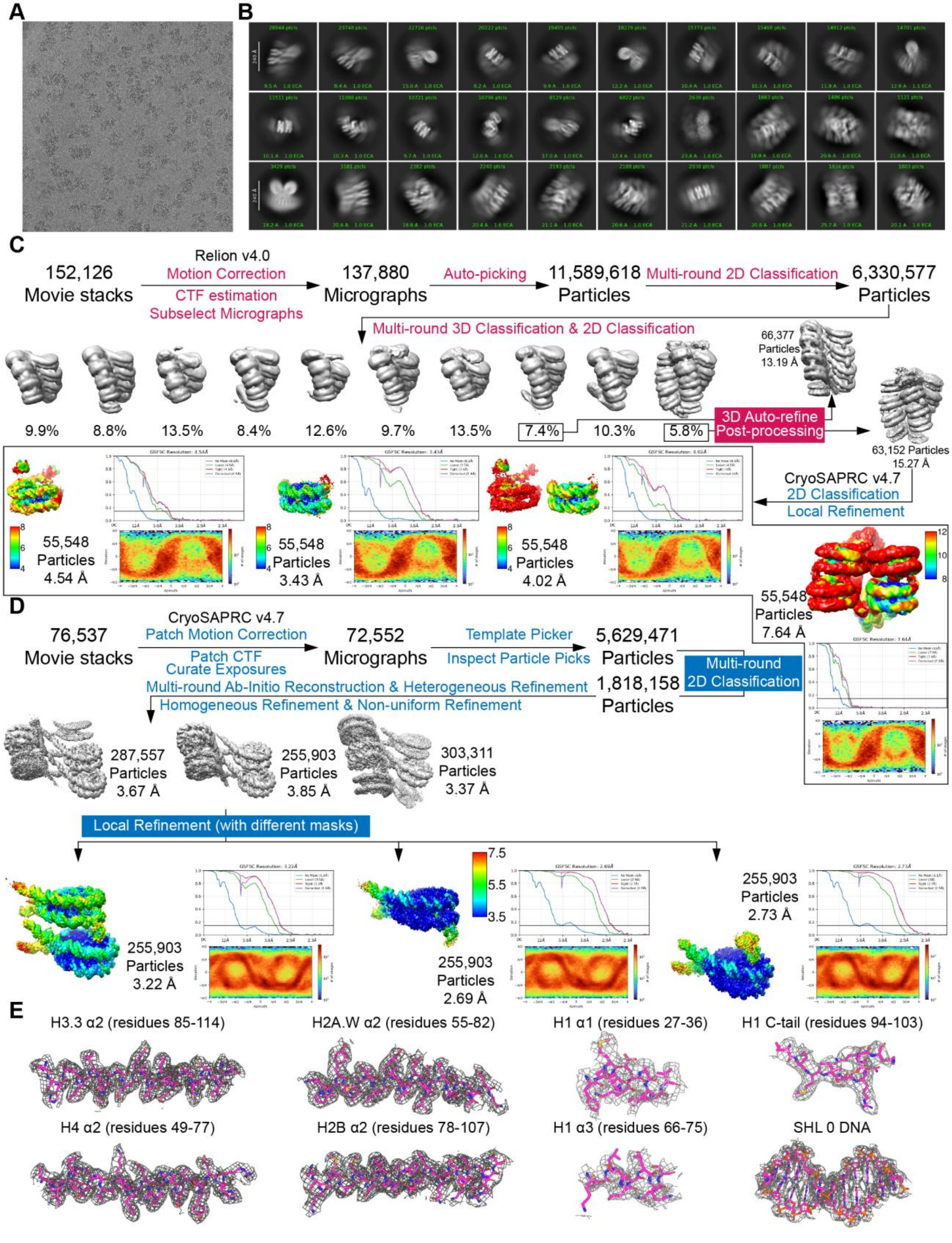
A
B
C
Relion v4.0
152,126 Movie stacks
Motion Correction
CTF estimation
Subselect Micrographs
137,880 Micrographs
Auto-picking
11,589,618 Particles
Multi-round 2D Classification
6,330,577 Particles
Multi-round 3D Classification & 2D Classification
66,377 Particles 13.19 Å
9.9%
8.8%
13.5%
8.4%
12.6%
9.7%
13.5%
7.4%
10.3%
5.8%
3D Auto-refine
Post-processing
63,152 Particles 15.27 Å
CryoSAPRC v4.7
2D Classification
Local Refinement
55,548 Particles 4.54 Å
55,548 Particles 3.43 Å
55,548 Particles 4.02 Å
55,548 Particles 7.64 Å
D
CryoSAPRC v4.7
76,537 Movie stacks
Patch Motion Correction
Patch CTF
Curate Exposures
72,552 Micrographs
Template Picker
Inspect Particle Picks
5,629,471 Particles
Multi-round 2D Classification
Multi-round Ab-Initio Reconstruction & Heterogeneous Refinement
Homogeneous Refinement & Non-uniform Refinement
1,818,158 Particles
287,557 Particles 3.67 Å
255,903 Particles 3.85 Å
303,311 Particles 3.37 Å
Local Refinement (with different masks)
255,903 Particles 3.22 Å
255,903 Particles 2.69 Å
255,903 Particles 2.73 Å
E
H3.3 α2 (residues 85-114)
H2A.W α2 (residues 55-82)
H1 α1 (residues 27-36)
H1 C-tail (residues 94-103)
H4 α2 (residues 49-77)
H2B α2 (residues 78-107)
H1 α3 (residues 66-75)
SHL 0 DNA

**Fig. S12. Cryo-EM data processing of the chromatin fiber containing H1.3, H2A.W, H2B, H3.3, H4 and 12-repeat 177-bp Widom 601 DNA (177_12_H2A.W_H3.3).**

(**A**) Representative cryo-EM micrograph collected using a Falcon 4i direct electron detector (4096 × 4096 pixels; pixel size, 0.959 Å).

(**B**) Selected reference-free two-dimensional (2D) class averages.

(**C**) Workflow of cryo-EM data processing performed in RELION v4.0.

(**D**) Workflow of cryo-EM data processing performed in cryoSPARC v4.7.

(**E**) Representative map-to-density fits for selected regions of the reconstructed chromatin fiber.

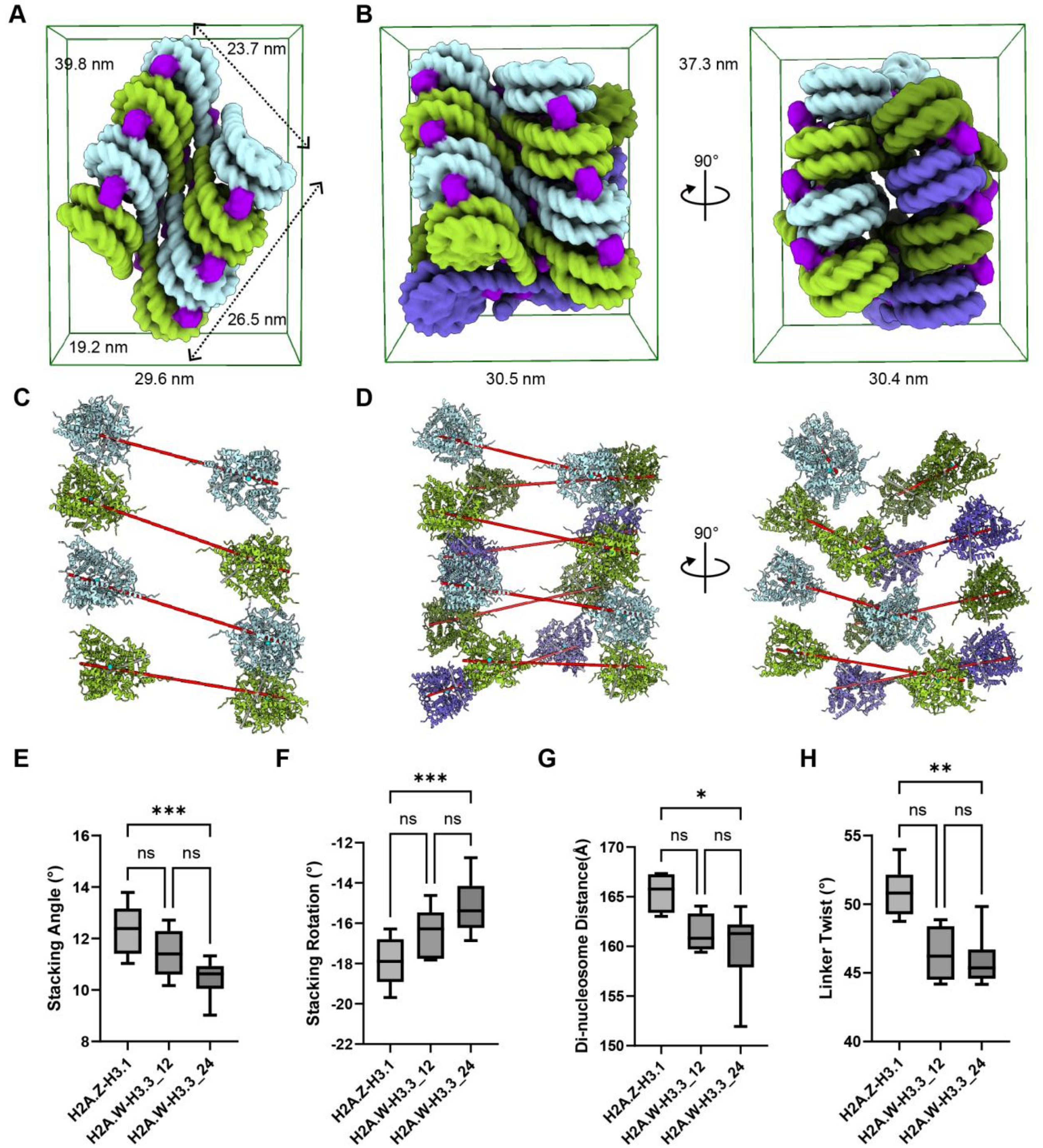


**Fig. S13. Quantitative analyses of nucleosome stacking parameters for Arabidopsis chromatin fiber and back-to-back fiber dimerization containing H2A.W and H3.3.**

(**A, B**) Dimensions of the cryo-EM density map of the Arabidopsis chromatin fiber 177_12_H2A.W_H3.3 (A) and back-to-back fiber dimerization (177_12_H2A.W_H3.3_di) (B) measured using the Blob tool in UCSF ChimeraX. Adjacent dinucleosome units are colored differently for clarity, with linker histone H1.3 highlighted in dark violet.

(**C, D**) Definitions of the mononucleosome axis and dinucleosome axis in the Arabidopsis chromatin fibers 177_12_H2A.W_H3.3 (C) and 177_12_H2A.W_H3.3_di (D). The dinucleosome axis was used to measure the inter dinucleosome angle between adjacent dinucleosome units.

(**E, F, G, H**) Box plots comparing stacking angles (E), stacking rotations (F), dinucleosome distances (G), linker twists (H), respectively, measured for the Arabidopsis chromatin fibers 177_12_H2A.Z (H2A.Z-H3.1), 177_12_H2A.W_H3.3 (H2A.W-H3.3_12) and 177_12_H2A.W_H3.3_di (H2A.W-H3.3_24). Statistical significance was assessed using the Kruskal-Wallis test followed by Dunn's multiple comparisons test. Asterisks indicate levels of significance (*$p < 0.05$; **$p < 0.01$; ***$p < 0.001$); ns, not significant.

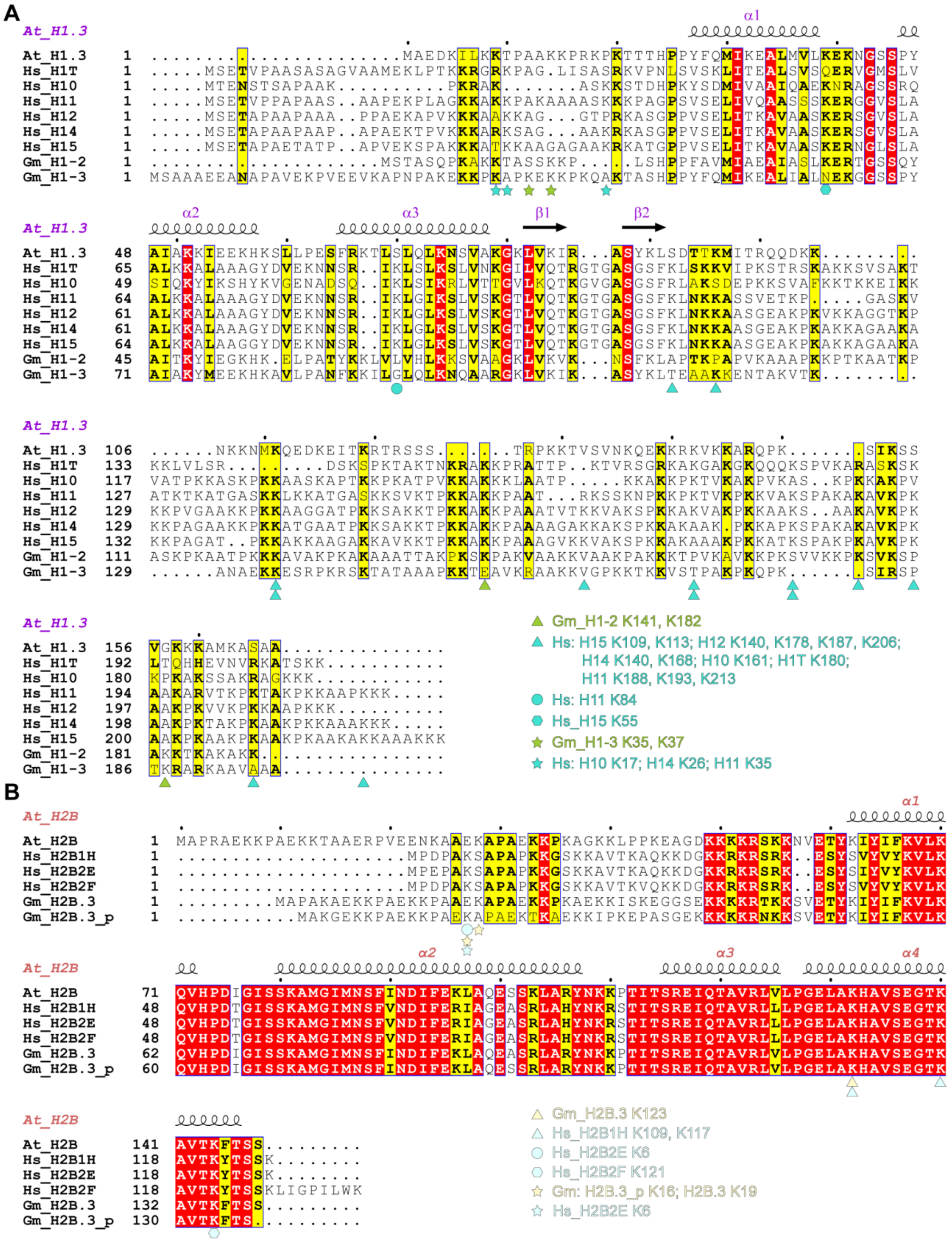
A
At_H1.3
Gm_H1-2 K141, K182
Hs: H15 K109, K113; H12 K140, K178, K187, K206; H14 K140, K168; H10 K161; H1T K180; H11 K188, K193, K213
Hs: H11 K84
Hs_H15 K55
Gm_H1-3 K35, K37
Hs: H10 K17; H14 K26; H11 K35
B
At_H2B
Gm_H2B.3 K123
Hs_H2B1H K109, K117
Hs_H2B2E K6
Hs_H2B2F K121
Gm: H2B.3_p K16; H2B.3 K19
Hs_H2B2E K6

**Fig. S14. Sequence alignments and the amino acids cross-linked between H2B and H1 histones identified in the soybean and human nuclei.**

(**A**) Sequence alignment of linker histone H1 highlighting all residues cross-linked with H2B, some of which are shown in Fig. 5, based on the published data of *in vivo* cross-linking mass spectrometry with soybean and human nuclei (*32, 33*). The UniProt accession numbers for Arabidopsis and human linker histone H1 are At_H1.3 (P94109), Hs_H1T (P22492), Hs_H10 (P07305), Hs_H11 (Q02539), Hs_H12 (P16403), Hs_H14 (P10412), and Hs_H15 (P16401). The NCBI RefSeq protein accession numbers for soybean histone H1 are Gm_H1-2 (predicted protein, XP_003536128.1) and Gm_H1-3 (curated protein, NP_001237870.2).

(**B**) Sequence alignment of histone H2B highlighting all residues cross-linked with H1, some of which are shown in Fig. 5, based on the published data of *in vivo* cross-linking mass spectrometry with soybean and human nuclei (*32, 33*). Distinct cross-linked regions are indicated using different colored symbols and fonts. Residue identities corresponding to the visible N- and C-terminal boundaries of the Arabidopsis H1 or H2B models, as well as their soybean or human counterparts, are shown in black. The UniProt accession numbers for Arabidopsis and human histone H2B proteins are At_H2B (Q9LQQ4), Hs_H2B1H (Q93079), Hs_H2B2E (Q16778), and Hs_H2B2F (Q5QNW6-2). The NCBI RefSeq protein accession numbers for soybean histone H2B are Gm_H2B.3 (predicted protein, XP_003542467.1) and Gm_H2B.3_p (curated protein, NP_001346041.1).

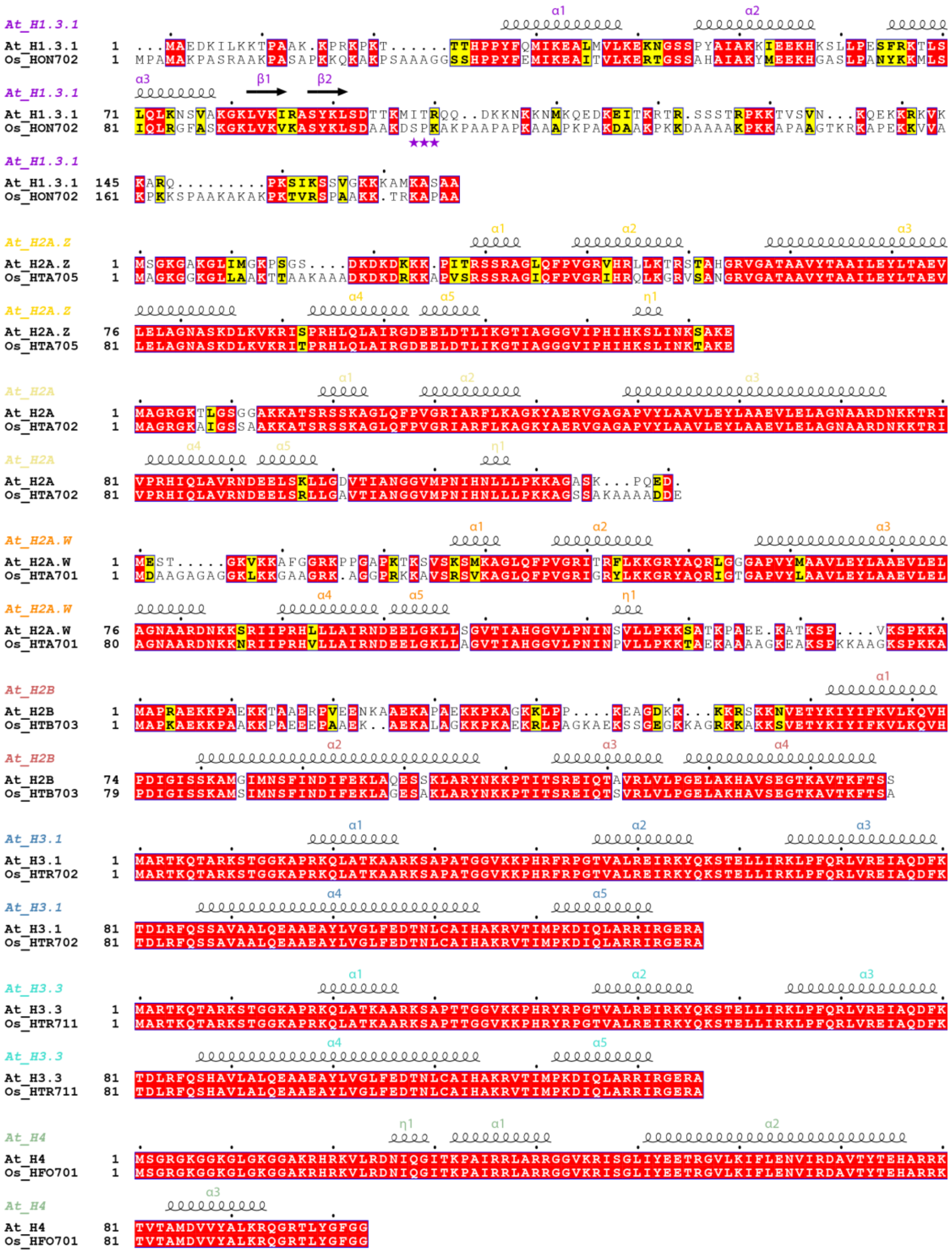

**Fig. S15. Sequence alignments of histones of *Arabidopsis thaliana* and *Oryza sativa*.**

Sequence alignments of linker histone H1.3, core histones and variants H2A, H2A.Z, H2A.W, H2B, H3.1, H3.3 and H4 between *Arabidopsis thaliana* and *Oryza sativa*. The asterisks indicate the key amino acid residues of H1.3 interacting with the nucleosomal acidic patch.

**Table S1. Cryo-EM data collection, refinement, and validation statistics for the H3.3 containing mononucleosomes.**

| | 152-H2A-H3.3 (PDB 9K40) | 152-H2A.Z-H3.3 (PDB 9K3Z) | 152-H2A.W-H3.3 (PDB 9K41) |
|---|---|---|---|
| **Data collection and processing** | | | |
| Microscope | Titan Krios | Titan Krios | Titan Krios |
| Camera | K3 Summit | K3 Summit | K3 Summit |
| Magnification | 64,000 | 81,000 | 81,000 |
| Voltage (kV) | 300 | 300 | 300 |
| Pixel size (Å) | 1.1 | 1.1 | 1.072 |
| Electron exposure ($e^{-}$/Å$^2$) | 50 | 50 | 50 |
| Exposure per frame ($e^{-}$/Å$^2$) | 1.25 | 1.25 | 1.25 |
| Number of frames collected | 40 | 40 | 40 |
| Defocus range (μm) | -0.9 to -1.8 | -0.9 to -1.8 | -0.9 to -1.8 |
| Symmetry imposed | C2 | C2 | C2 |
| Micrographs (no.) | 5,202 | 3,082 | 1,637 |
| Initial particle images (no.) | 3,670,476 | 4,623,825 | 1,649,336 |
| Final particle images (no.) | 149,713 | 240,998 | 165,777 |
| Final reconstruction package | cryoSPARC | cryoSPARC | cryoSPARC |
| C1 Map resolution (Å) | 3.47 | 2.9 | 2.97 |
| C1 Map EMDB ID | EMD-62043 | EMD-62045 | EMD-62047 |
| C2 Map resolution (Å) | 3.22 | 2.71 | 2.85 |
| C2 Map EMDB ID | EMD-62044 | EMD-62046 | EMD-62048 |
| FSC threshold | 0.143 | 0.143 | 0.143 |
| **Refinement** | | | |
| Refinement package | Phenix | Phenix | Phenix |
| Initial model used (PDB) | 7OHC | 7OHC | 7OHC |
| Map sharpening B factor (Å$^2$) | -103.2 | -99.6 | -104.8 |
| **Model composition** | | | |
| Non-hydrogen atoms | 11,840 | 11,830 | 11,844 |
| Protein residues | 746 | 744 | 746 |
| Nucleotide residues | 290 | 290 | 290 |
| Ligands | 0 | 0 | 0 |
| **R.m.s. deviations** | | | |
| Bond lengths (Å) | 0.004 | 0.004 | 0.004 |
| Bond angles (°) | 0.749 | 0.767 | 0.758 |
| **Validation** | | | |
| MolProbity score | 1.41 | 1.26 | 1.34 |
| Clash score | 7.4 | 4.97 | 6.22 |
| Rotamer outliers (%) | 0.16 | 0.96 | 0.8 |
| Cβ outliers (%) | 0 | 0 | 0 |
| CaBLAM outliers (%) | 0.84 | 0.98 | 0.56 |
| **Overall correlation coefficients** | | | |
| CC (mask) | 0.84 | 0.76 | 0.8 |
| CC (box) | 0.83 | 0.69 | 0.72 |
| CC (peaks) | 0.78 | 0.62 | 0.66 |
| CC (volume) | 0.82 | 0.75 | 0.79 |
| **Ramachandran plot** | | | |
| Favored (%) | 98.77 | 98.63 | 98.49 |
| Allowed (%) | 1.23 | 1.37 | 1.51 |
| Disallowed (%) | 0 | 0 | 0 |

**Table S2. Cryo-EM data collection, refinement, and validation statistics for the 177_12_H2A.Z chromatin fiber.**

| | | | | 177_12_H2A.Z | | | |
|---|---|---|---|---|---|---|---|
| **Data collection and processing** | | | | | | | |
| Microscope | | | | Titan Krios | | | |
| Camera | | | | Falcon 4i | | | |
| Magnification | | | | 130,000 | | | |
| Voltage (kV) | | | | 300 | | | |
| Pixel size (Å) | | | | 0.959 | | | |
| Electron exposure (e⁻/Å²) | | | | 40 | | | |
| Defocus range (μm) | | | | -0.6 to -1.6 | | | |
| Micrographs (no.) | | | | 76,537 | | | |
| Initial particle images (no.) | | | | 6,025,056 | | | |
| Symmetry imposed | | | | C1 | | | |
| Final particle images (no.) | 50,171 | 294,094 | 36,592 | 30,131 | 13,924 | 99,433 | |
| Final reconstruction package | RELION | | | CryoSPARC | | | Composite map |
| Map resolution (Å) | 10.42 | 3.74 | 3.44 | 3.48 | 3.21 | 2.87 | |
| FSC threshold | 0.143 | 0.143 | 0.143 | 0.143 | 0.143 | 0.143 | N/A |
| Map EMDB ID (EMD-) | 67700 | 67701 | 67702 | 67703 | 67704 | 67705 | 67706 |
| PDB ID | | | | | | 21IU | 21IV |
| **Refinement** | | | | | | | |
| Refinement package | | | | | | Phenix | Phenix |
| Initial model used (PDB) | | | | | | 9K3Z | 9K3Z |
| **Model composition** | | | | | | | |
| Non-hydrogen atoms | | | | | | 14,906 | 29,245 |
| Protein residues | | | | | | 967 | 1,863 |
| Nucleotide residues | | | | | | 354 | 708 |
| Ligands | | | | | | 0 | 0 |
| **R.m.s. deviations** | | | | | | | |
| Bond lengths (Å) | | | | | | 0.009 | 0.009 |
| Bond angles (°) | | | | | | 0.964 | 0.965 |
| **Validation** | | | | | | | |
| MolProbity score | | | | | | 2.07 | 2.03 |
| Clash score | | | | | | 8.33 | 8.99 |
| Rotamer outliers (%) | | | | | | 4.02 | 3.43 |
| Cβ outliers (%) | | | | | | 0 | 0 |
| CaBLAM outliers (%) | | | | | | 1.30 | 1.01 |
| **Overall correlation coefficients** | | | | | | | |
| CC (mask) | | | | | | 0.81 | 0.74 |
| CC (box) | | | | | | 0.72 | 0.72 |
| CC (peaks) | | | | | | 0.67 | 0.64 |
| CC (volume) | | | | | | 0.81 | 0.75 |
| **Ramachandran plot** | | | | | | | |
| Favored (%) | | | | | | 97.14 | 97.20 |
| Allowed (%) | | | | | | 2.33 | 2.47 |
| Disallowed (%) | | | | | | 0.53 | 0.33 |

**Table S3. The stacking distances, stacking angles, and rotational angles between N and N+2 nucleosomes within four types of chromatin fibers including H5-chromatin fiber (PDB: 8XJV; EMD-38407), and Arabidopsis chromatin fibers determined in this study: 177_12_H2A.Z (Z31, EMD-67700), 177_12_H2A.W_H3.3 (W33-12, EMD-68924) and 177_12_H2A.W_H3.3_di (W33-24, EMD-67724).**

| | **8XJV** | **Z31** | **W33-12** | **W33-24-1** | **W33-24-2** |
|---|---|---|---|---|---|
| **Stacking Distance (Å)** | | | | | |
| 1 - 3 | 59.361 | 73.241 | 74.882 | 76.47 | 76.253 |
| 2 - 4 | 57.355 | 73.674 | 75.363 | 78.931 | 74.505 |
| 3 - 5 | 65.374 | 73.818 | 74.829 | 75.348 | 74.717 |
| 4 - 6 | 64.416 | 73.953 | 75.668 | 74.874 | 74.744 |
| 5 - 7 | 58.559 | 73.982 | 74.814 | 75.478 | 74.244 |
| 6 - 8 | 57.738 | 73.755 | 75.072 | 75.019 | 77.58 |
| 7 - 9 | 63.581 | 73.895 | | | |
| 8 - 10 | 61.614 | 73.29 | | | |
| 9 - 11 | 57.956 | 72.863 | | | |
| 10 - 12 | 58.646 | 72.98 | | | |
| Average Value | 60.46 | 73.5451 | 75.10467 | 76.02 | 75.3405 |
| **Stacking Angle (°)** | | | | | |
| 1 - 3 | 21.663 | 11.032 | 10.172 | 9.026 | 9.982 |
| 2 - 4 | 20.589 | 13.789 | 12.152 | 11.325 | 10.786 |
| 3 - 5 | 5.592 | 12.427 | 11.373 | 10.401 | 10.934 |
| 4 - 6 | 20.243 | 12.969 | 11.431 | 10.831 | 10.246 |
| 5 - 7 | 7.202 | 12.358 | 12.716 | 10.918 | 11.307 |
| 6 - 8 | 3.49 | 12.176 | 10.745 | 10.473 | 9.38 |
| 7 - 9 | 6.296 | 12.801 | | | |
| 8 - 10 | 6.939 | 11.479 | | | |
| 9 - 11 | 10.132 | 13.758 | | | |
| 10 - 12 | 7.135 | 11.19 | | | |
| Average Value | 10.9281 | 12.3979 | 11.4315 | 10.49567 | 10.43917 |
| **Stacking Rotation (°)** | | | | | |
| 1 - 3 | -9.548 | -16.284 | -14.618 | -12.745 | -13.987 |
| 2 - 4 | -24.662 | -19.687 | -17.741 | -15.824 | -16.865 |
| 3 - 5 | -22.206 | -17.774 | -16.179 | -14.81 | -16.037 |
| 4 - 6 | -16.89 | -18.715 | -16.373 | -16.029 | -14.931 |
| 5 - 7 | -14.229 | -18.026 | -17.829 | -16.289 | -16.525 |
| 6 - 8 | -15.461 | -17.456 | -15.743 | -14.631 | -13.503 |
| 7 - 9 | -22.848 | -18.255 | | | |
| 8 - 10 | -25.668 | -16.882 | | | |
| 9 - 11 | -17.29 | -19.518 | | | |
| 10 - 12 | -17.036 | -16.521 | | | |
| Average Value | -18.5838 | -17.9118 | -16.4138 | -15.0547 | -15.308 |

**Table S4. Dinucleosome distances, opening angles, linker twist angles, and dinucleosome rotation angles within four types of chromatin fibers, including the previously reported animal H5-chromatin fiber (PDB: 8XJV; EMD-38407) and Arabidopsis chromatin fibers determined in this study: 177_12_H2A.Z (Z31, EMD-67700), 177_12_H2A.W_H3.3 (W33-12, EMD-68924) and 177_12_H2A.W_H3.3_di (W33-24, EMD-67724).**

| | 8XJV | Z31 | W33-12 | W33-24-1 | W33-24-2 |
|---|---|---|---|---|---|
| **Dinucleosome distance (Å)** | | | | | |
| 1 - 2 | 186.065 | 163.02 | 164.061 | 151.944 | 159.594 |
| 3 - 4 | 189.116 | 165.317 | 161.045 | 161.578 | 161.739 |
| 5 - 6 | 199.768 | 166.23 | 159.411 | 161.014 | 162.372 |
| 7 - 8 | 197.313 | 167.235 | 160.605 | 164.031 | 157.333 |
| 9 - 10 | 191.172 | 167.344 | | | |
| 11 - 12 | 176.981 | 163.52 | | | |
| Average Value | 190.0692 | 165.4443 | 161.2805 | 159.6418 | 160.2595 |
| **Opening Angle (°)** | | | | | |
| 1 - 2 | 35.209 | 63.977 | 65.244 | 74.373 | 66.052 |
| 3 - 4 | 36.643 | 63.687 | 63.618 | 63.162 | 62.557 |
| 5 - 6 | 39.619 | 63.684 | 63.48 | 62.942 | 63.088 |
| 7 - 8 | 44.351 | 64.758 | 63.452 | 63.665 | 67.657 |
| 9 - 10 | 54.469 | 64.764 | | | |
| 11 - 12 | 49.718 | 66.377 | | | |
| Average Value | 43.33483 | 64.54117 | 63.9485 | 66.0355 | 64.8385 |
| **Linker Twist (°)** | | | | | |
| 1 - 2 | -176.373 | 51.542 | 48.871 | 45.278 | 47.083 |
| 3 - 4 | 158.648 | 50.134 | 45.484 | 44.175 | 45.217 |
| 5 - 6 | -151.375 | 49.436 | 44.18 | 44.363 | 45.594 |
| 7 - 8 | -166.851 | 48.755 | 46.95 | 45.467 | 49.837 |
| 9 - 10 | 118.093 | 51.498 | | | |
| 11 - 12 | 173.326 | 53.982 | | | |
| Average Value | -7.422 | 50.89117 | 46.37125 | 44.82075 | 46.93275 |
| **Dinucleosome rotation Angle (°)** | | | | | |
| 12 - 34 | 34.856 | 25.159 | 18.855 | 15.982 | 17.205 |
| 34 - 56 | 20.112 | 23.029 | 18.265 | 18.357 | 18.622 |
| 56 - 78 | 30.73 | 22.159 | 19.84 | 18.019 | 16.503 |
| 78 - 910 | 13.4 | 22.287 | | | |
| 1011 - 1112 | 33.303 | 25.117 | | | |
| Average Value | 26.4802 | 23.5502 | 18.98667 | 17.45267 | 17.44333 |

**Table S5. Cryo-EM data collection, refinement, and validation statistics for the 177_12_H2A chromatin fiber.**

| | | | 177_12_H2A | | |
|---|---|---|---|---|---|
| **Data collection and processing** | | | | | |
| Microscope | | | Titan Krios | | |
| Camera | | | Falcon 4i | | |
| Magnification | | | 130,000 | | |
| Voltage (kV) | | | 300 | | |
| Pixel size (Å) | | | 0.959 | | |
| Electron exposure ($e^{-}/Å^{2}$) | | | 40 | | |
| Defocus range (μm) | | | -0.6 to -1.6 | | |
| Micrographs (no.) | | | 37,290 | | |
| Initial particle images (no.) | | | 3,217,978 | | |
| Symmetry imposed | | | C1 | | |
| Final particle images (no.) | 205,012 | 200,318 | 200,318 | 200,318 | 200,318 |
| Final reconstruction package | | | CryoSPARC | | |
| Map resolution (Å) | 4.22 | 4.04 | 2.80 | 2.81 | 3.74 |
| FSC threshold | | | 0.143 | | |
| Map EMDB ID (EMD-) | 67707 | 67708 | 67709 | 67710 | 67711 |
| PDB ID | | | | | 21IW |
| **Refinement** | | | | | |
| Refinement package | | | | | Phenix |
| Initial model used (PDB) | | | | | 9K3Z |
| **Model composition** | | | | | |
| Non-hydrogen atoms | | | | | 28,036 |
| Protein residues | | | | | 1,714 |
| Nucleotide residues | | | | | 708 |
| Ligands | | | | | 0 |
| **R.m.s. deviations** | | | | | |
| Bond lengths (Å) | | | | | 0.004 |
| Bond angles (°) | | | | | 0.768 |
| **Validation** | | | | | |
| MolProbity score | | | | | 2.06 |
| Clash score | | | | | 9.87 |
| Rotamer outliers (%) | | | | | 2.38 |
| Cβ outliers (%) | | | | | 0 |
| CaBLAM outliers (%) | | | | | 1.34 |
| **Overall correlation coefficients** | | | | | |
| CC (mask) | | | | | 0.80 |
| CC (box) | | | | | 0.86 |
| CC (peaks) | | | | | 0.74 |
| CC (volume) | | | | | 0.79 |
| **Ramachandran plot** | | | | | |
| Favored (%) | | | | | 96.24 |
| Allowed (%) | | | | | 3.70 |
| Disallowed (%) | | | | | 0.06 |

**Table S6. Cryo-EM data collection, refinement, and validation statistics for the 177_12_H2A.W chromatin fiber.**

| | | | 177_12_H2A.W | | |
|---|---|---|---|---|---|
| **Data collection and processing** | | | | | |
| Microscope | | | Titan Krios | | |
| Camera | | | Falcon 4i | | |
| Magnification | | | 130,000 | | |
| Voltage (kV) | | | 300 | | |
| Pixel size (Å) | | | 0.959 | | |
| Electron exposure (e⁻/Å²) | | | 40 | | |
| Defocus range (μm) | | | -0.6 to -1.6 | | |
| Micrographs (no.) | | | 84,073 | | |
| Initial particle images (no.) | | | 7,209,456 | | |
| Symmetry imposed | | | C1 | | |
| Final particle images (no.) | 203290 | 214595 | 214595 | 214595 | 214595 |
| Final reconstruction package | | | CryoSPARC | | |
| Map resolution (Å) | 3.79 | 3.67 | 2.72 | 2.89 | 3.05 |
| FSC threshold | | | 0.143 | | |
| Map EMDB ID (EMD-) | 67712 | 67713 | 67714 | 67715 | 67716 |
| PDB ID | | | | | 21IX |
| **Refinement** | | | | | |
| Refinement package | | | | | Phenix |
| Initial model used (PDB) | | | | | 9K3Z |
| **Model composition** | | | | | |
| Non-hydrogen atoms | | | | | 27,983 |
| Protein residues | | | | | 1,707 |
| Nucleotide residues | | | | | 708 |
| Ligands | | | | | 0 |
| **R.m.s. deviations** | | | | | |
| Bond lengths (Å) | | | | | 0.004 |
| Bond angles (°) | | | | | 0.758 |
| **Validation** | | | | | |
| MolProbity score | | | | | 1.65 |
| Clash score | | | | | 9.72 |
| Rotamer outliers (%) | | | | | 0.70 |
| Cβ outliers (%) | | | | | 0 |
| CaBLAM outliers (%) | | | | | 0.98 |
| **Overall correlation coefficients** | | | | | |
| CC (mask) | | | | | 0.65 |
| CC (box) | | | | | 0.75 |
| CC (peaks) | | | | | 0.58 |
| CC (volume) | | | | | 0.64 |
| **Ramachandran plot** | | | | | |
| Favored (%) | | | | | 97.24 |
| Allowed (%) | | | | | 2.64 |
| Disallowed (%) | | | | | 0.12 |

**Table S7. Cryo-EM data collection, refinement, and validation statistics for the 177_12_H2A.Z_H5 chromatin fiber.**

| | 177_12_H2A.Z-H5 | | | |
|---|---|---|---|---|
| **Data collection and processing** | | | | |
| Microscope | Titan Krios | | | |
| Camera | Falcon 4i | | | |
| Magnification | 130,000 | | | |
| Voltage (kV) | 300 | | | |
| Pixel size (Å) | 0.959 | | | |
| Electron exposure ($e^-$/Å$^2$) | 40 | | | |
| Defocus range (μm) | -0.6 to -1.6 | | | |
| Micrographs (no.) | 31,432 | | | |
| Initial particle images (no.) | 2,609,081 | | | |
| Symmetry imposed | C1 | | | |
| Final particle images (no.) | 111,187 | 111,187 | 111,187 | 37,844 |
| Final reconstruction package | CryoSPARC | | | |
| Map resolution (Å) | 3.07 | 2.96 | 3.92 | 3.85 |
| FSC threshold | 0.143 | | | |
| Map EMDB ID (EMD-) | 67717 | 67718 | 67719 | 67720 |
| PDB ID | | | 21IY | 21IZ |
| **Refinement** | | | | |
| Refinement package | | | Phenix | Phenix |
| Initial model used (PDB) | | | 9K3Z | 9K3Z |
| **Model composition** | | | | |
| Non-hydrogen atoms | | | 27,366 | 13,710 |
| Protein residues | | | 1,638 | 822 |
| Nucleotide residues | | | 708 | 354 |
| Ligands | | | 0 | 0 |
| **R.m.s. deviations** | | | | |
| Bond lengths (Å) | | | 0.008 | 0.008 |
| Bond angles (°) | | | 0.944 | 0.950 |
| **Validation** | | | | |
| MolProbity score | | | 1.61 | 1.36 |
| Clash score | | | 7.20 | 6.29 |
| Rotamer outliers (%) | | | 1.85 | 1.03 |
| Cβ outliers (%) | | | 0 | 0 |
| CaBLAM outliers (%) | | | 1.34 | 1.15 |
| **Overall correlation coefficients** | | | | |
| CC (mask) | | | 0.56 | 0.63 |
| CC (box) | | | 0.74 | 0.75 |
| CC (peaks) | | | 0.51 | 0.59 |
| CC (volume) | | | 0.57 | 0.64 |
| **Ramachandran plot** | | | | |
| Favored (%) | | | 97.94 | 98.76 |
| Allowed (%) | | | 1.81 | 1.12 |
| Disallowed (%) | | | 0.25 | 0.12 |

**Table S8. Cryo-EM data collection, refinement, and validation statistics for the 177_12_H2A.W_H3.3 chromatin fiber.**

| | | | | | | | | | |
|---|---|---|---|---|---|---|---|---|---|
| | | | | | **177_12_H2A.W_H3.3** | | | | |
| **Data collection and processing** | | | | | | | | | |
| Microscope | | | | | Titan Krios | | | | |
| Camera | | | | | Falcon 4i | | | | |
| Magnification | | | | | 130,000 | | | | |
| Voltage (kV) | | | | | 300 | | | | |
| Pixel size (Å) | | | | | 0.959 | | | | |
| Electron exposure (e⁻/Å²) | | | | | 40 | | | | |
| Defocus range (μm) | | | | | -0.6 to -1.6 | | | | |
| Micrographs (no.) | | | | | 152,126 | | | | |
| Initial particle images (no.) | | | | | 11,589,618 | | | | |
| Symmetry imposed | | | | | C1 | | | | |
| Final particle images (no.) | 255,903 | 255,903 | 255,903 | 63,152 | 55,548 | 55,548 | 55,548 | 55,548 | 66,377 |
| Final reconstruction package | | CryoSPARC | | RELION | | CryoSPARC | | | RELION |
| Map resolution (Å) | 2.73 | 2.69 | 3.22 | 15.27 | 7.64 | 3.43 | 4.54 | 4.02 | 13.19 |
| FSC threshold | | | | | 0.143 | | | | |
| Map EMDB ID (EMD-) | 67721 | 67722 | 67723 | 67724 | 67725 | 67726 | 67727 | 67728 | 68924 |
| PDB ID | | | 21JA | | | | | 21JB | |
| **Refinement** | | | | | | | | | |
| Refinement package | | | Phenix | | | | | Phenix | |
| Initial model used (PDB) | | | 9K3Z | | | | | 9K3Z | |
| **Model composition** | | | | | | | | | |
| Non-hydrogen atoms | | | 28,008 | | | | | 27,346 | |
| Protein residues | | | 1,706 | | | | | 1,624 | |
| Nucleotide residues | | | 708 | | | | | 708 | |
| Ligands | | | 0 | | | | | 0 | |
| **R.m.s. deviations** | | | | | | | | | |
| Bond lengths (Å) | | | 0.005 | | | | | 0.005 | |
| Bond angles (°) | | | 0.757 | | | | | 0.750 | |
| **Validation** | | | | | | | | | |
| MolProbity score | | | 1.75 | | | | | 1.65 | |
| Clash score | | | 9.59 | | | | | 9.84 | |
| Rotamer outliers (%) | | | 1.18 | | | | | 0.37 | |
| Cβ outliers (%) | | | 0 | | | | | 0 | |
| CaBLAM outliers (%) | | | 0.92 | | | | | 0.71 | |
| **Overall correlation coefficients** | | | | | | | | | |
| CC (mask) | | | 0.78 | | | | | 0.62 | |
| CC (box) | | | 0.81 | | | | | 0.62 | |
| CC (peaks) | | | 0.71 | | | | | 0.44 | |
| CC (volume) | | | 0.77 | | | | | 0.61 | |
| **Ramachandran plot** | | | | | | | | | |
| Favored (%) | | | 96.88 | | | | | 97.29 | |
| Allowed (%) | | | 2.94 | | | | | 2.71 | |
| Disallowed (%) | | | 0.18 | | | | | 0 | |

**Table S9. Selected cross-linking mass spectrometry data for linker histone H1 and core histone H2B of *Homo sapiens* and *Glycine max*, derived from the published datasets (*32, 33*) and used in Fig. 5 and Fig. S14.**

| Species | Pro1 | PepSeq1 | Pro1 ID | Pro2 | PepSeq2 | Pro2 ID | Pro1-Pro2 sites |
|---|---|---|---|---|---|---|---|
| **(1) C-terminal region of H1 & C-terminal α4 helix of H2B** | | | | | | | |
| *Homo sapiens* | H15 | GTGASGSFKLNK | P16401 | H2B1H | LLLPGELAKHAVSEGTK | Q93079 | 109-109 |
| *Homo sapiens* | H15 | KAASGEAKPK | P16401 | H2B1H | LLLPGELAKHAVSEGTK | Q93079 | 113-109 |
| *Homo sapiens* | H14 | KATGAATPK | P10412 | H2B1H | LLLPGELAKHAVSEGTK | Q93079 | 140-109 |
| *Homo sapiens* | H12 | KAAGGATPK | P16403 | H2B1H | LLLPGELAKHAVSEGTK | Q93079 | 140-109 |
| *Homo sapiens* | H10 | KPKTVK | P07305 | H2B1H | LLLPGELAKHAVSEGTK | Q93079 | 161-109 |
| *Homo sapiens* | H14 | KPAAAAGAKK | P10412 | H2B1H | LLLPGELAKHAVSEGTK | Q93079 | 168-109 |
| *Homo sapiens* | H12 | AKVAKPK | P16403 | H2B1H | LLLPGELAKHAVSEGTK | Q93079 | 178-109 |
| *Homo sapiens* | H1T | QQQKSPVKAR | P22492 | H2B1H | LLLPGELAKHAVSEGTK | Q93079 | 180-109 |
| *Homo sapiens* | H12 | AAKSAAK | P16403 | H2B1H | LLLPGELAKHAVSEGTK | Q93079 | 187-109 |
| *Homo sapiens* | H11 | AKAVKPK | Q02539 | H2B1H | LLLPGELAKHAVSEGTK | Q93079 | 188-109 |
| *Homo sapiens* | H11 | AVKPKAAK | Q02539 | H2B1H | LLLPGELAKHAVSEGTK | Q93079 | 193-109 |
| *Homo sapiens* | H12 | VVKPKK | P16403 | H2B1H | LLLPGELAKHAVSEGTK | Q93079 | 206-109 |
| *Homo sapiens* | H11 | AAPKK | Q02539 | H2B1H | LLLPGELAKHAVSEGTK | Q93079 | 213-109 |
| *Homo sapiens* | H11 | AAPKKK | Q02539 | H2B1H | HAVSEGTKAVTK | Q93079 | 213-117 |
| *Glycine max* | H1-2 | SKPAK | XP_003536128.1 | H2B.3 | LVLPGELAKHAVSEGTK | XP_003542467.1 | 141-123 |
| *Glycine max* | H1-2 | SPAKK | XP_003536128.1 | H2B.3 | LVLPGELAKHAVSEGTK | XP_003542467.1 | 182-123 |
| *Glycine max* | H1-1 | KAAPAAKPAAK | XP_003536890.1 | H2B.3 | LVLPGELAKHAVSEGTK | XP_003542467.1 | 253-123 |
| *Glycine max* | H1-4 | AAAAKPAAKPK | XP_003518887.1 | H2B.3 | LVLPGELAKHAVSEGTK | XP_003542467.1 | 235-123 |
| **(2) Helix α3 of H1 & N-terminal region of H2B** | | | | | | | |

| | | | | | | | |
|---|---|---|---|---|---|---|---|
| *Homo sapiens* | H11 | IKLGIK | Q02539 | H2B2E | PEPAKSAPAPK | Q16778 | 84-6 |
| **(3) Regions spanning helices α1 and α2 of H1.5 & C-terminal helix α4 of H2B** | | | | | | | |
| *Homo sapiens* | H15 | AVAASKER | P16401 | H2B2F | AVTKYTSSK | Q5QNW6-2 | 55-121 |
| **(4) N-terminal regions of H1 & H2B** | | | | | | | |
| *Homo sapiens* | H10 | AKASK | P07305 | H2B2E | PEPAKSAPAPK | Q16778 | 17-6 |
| *Homo sapiens* | H14 | KSAGAAK | P10412 | H2B2E | PEPAKSAPAPK | Q16778 | 26-6 |
| *Homo sapiens* | H11 | AAAASKK | Q02539 | H2B2E | PEPAKSAPAPK | Q16778 | 35-6 |
| *Glycine max* | H1-3 | APKEK | NP_001237870.2 | H2B_p | KPAEKAPAEK | NP_001346041.1 | 35-16 |
| *Glycine max* | H1-3 | APKEKK | NP_001237870.2 | H2B.3 | KPAAEKAPAEK | XP_003542467.1 | 37-19 |
| *Glycine max* | H1-3 | APKEK | NP_001237870.2 | H2B | STVGDKAPAEK | XP_003539691.2 | 35-25 |
